# Cell-matrix mechanosensing and cellular metabolic demand are linked through SKT and mTORC2

**DOI:** 10.64898/2026.02.03.702586

**Authors:** Milan Collins, Lorna Young, Emily Goodall, Beth Hammond, Daniel Newman, Paul Atherton, Patrick Caswell, Tobias Zech

## Abstract

Integrin-based adhesion complexes mediate cell adhesion to the extracellular matrix and enable the cell to interpret and respond to both biochemical and mechanical cues. Such cues can affect a cells’ metabolic rate; however, how mechanical signals are converted into metabolic rate changes is not clear. We identified an uncharacterised protein; Sickle Tail Protein Homolog (KIAA1217), ‘SKT’ to be enriched in cell-matrix adhesion complexes in stiff microenvironments. Low SKT expression correlates with an improved prognosis in pancreatic ductal adenocarcinoma (PDAC), suggesting an important role for SKT in extracellular matrix dependent tumour progression. Here, we show that SKT interacts with the mechanistic target of rapamycin complex 2 (mTORC2), a pivotal signalling complex in glucose metabolism, cell growth, and survival. SKT recruits mTORC2 to cell-matrix adhesions in a mechanoresponsive manner. SKT mediated mTORC2 signalling from adhesions is required for maintaining glycolytic flux and control of adhesion dynamics. Our findings show that SKT serves as a rheostat that controls metabolic adaptation of cells to their matrix microenvironment. Collectively, our research provides insights into the molecular mechanisms and interplay between cell adhesion and metabolic signalling in complex and stiff tumour microenvironments. The novel functions identified for SKT in cell-matrix adhesions, mTORC2 signalling and glycolysis unveils a signalling axis between the tumour microenvironment and cellular metabolism that are required for PDAC growth and invasion.

## Introduction

Integrin-based adhesion complexes (IACs) are mechanosensitive signalling complexes whose formation and composition dynamically alter in response to biochemical and mechanical cues under varying physiological conditions^1–4^. Within IACs, activated Integrins act as scaffolds for the recruitment and binding of intracellular signalling and adapter proteins including actin and microtubule binding proteins, kinases, phosphatases, and small GTPases^5–7^. Collectively, the spatiotemporal interactions of IAC proteins contribute to the physiological function of cell adhesions, regulating both signal and mechano-transduction, and physically linking the cellular actomyosin cytoskeleton with the extracellular matrix (ECM)^4, 7–13^.

Previous studies have implicated mTOR (a highly conserved serine/threonine protein kinase that functions as a central regulator of cell growth and metabolism^14, 15^) activity in the regulation of cell adhesions^16^, and suggested mechanically induced recruitment of Akt (the downstream target of mTOR) to IACs^17, 18^. Furthermore, ECM substrate stiffness has been linked to intracellular energy regulation^19–21^. Therefore, it is a distinct possibility that key components of IACs may contribute to the regulation of cellular metabolism^22–24^, and provide a way for cells to adjust their metabolic plasticity in response to changes in matrix environment. In solid tumours such as Pancreatic Ductal Adenocarcinoma (PDAC), ECM remodeling stiffens the tumour microenvironment and metabolic changes that promote cancer cell survival and proliferation occur^25–29^. However, the underlying molecular mechanisms connecting ECM sensing to metabolic signalling remain poorly defined.

Here, we identify Sickle Tail Protein Homolog (SKT), as a novel IAC protein that is preferentially enriched on stiff microenvironments. We show that SKT is an mTORC2 interaction partner that regulates mTORC2 recruitment to adhesion complexes in response to increased substrate stiffness. This novel adhesion signalling component is required for adhesion turnover and cytoskeletal reorganisation on stiff matrices. SKT mediates localised mTORC2 and Akt activation at IACs to drive adjustments of glucose metabolism levels in response to increased cellular energy demands in stiffer ECM environments. Together, this reveals a tight coupling of IAC mediated mechanoresponses and metabolic rates that dictate metabolic signalling for tumour progression and maintenance.

## Results

### SKT enriches at adhesions on stiff microenvironments

Currently, there is no published data describing the localisation of SKT. To validate our previous BioID2 screen showing preferential enrichment of SKT at IACs on stiff microenvironments^28^ and and investigate whether SKT does in fact localise to IACs, endogenous SKT enrichment was quantified on soft (1.5 kPa) and stiff microenvironments (Glass, 1 GPa). Immunofluorescence revealed a significant increase in SKT intensity at IACs relative to cytoplasmic intensity when cells were plated on glass, in comparison to the soft 1.5 kPa substrates (**Fig. 1 a, b)**. To identify recruitment relative to established adhesion markers we transfected cells with GFP-SKT, and mAPPLE-paxillin to visualise IACs (**Fig. 1c**). Cells were seeded onto imaging dishes coated with fibronectin/collagen and time lapse microscopy was conducted to capture adhesion dynamics. GFP-SKT displayed distinct co-localisation with mAPPLE-paxillin during the formation of nascent adhesion complexes and remained co-localised throughout IAC lifetime (assembly to disassembly) (**Fig. 1c**). These imaging experiments demonstrate that SKT localises to integrin adhesion complexes in a stiffness-dependent manner.

**Figure 1.**
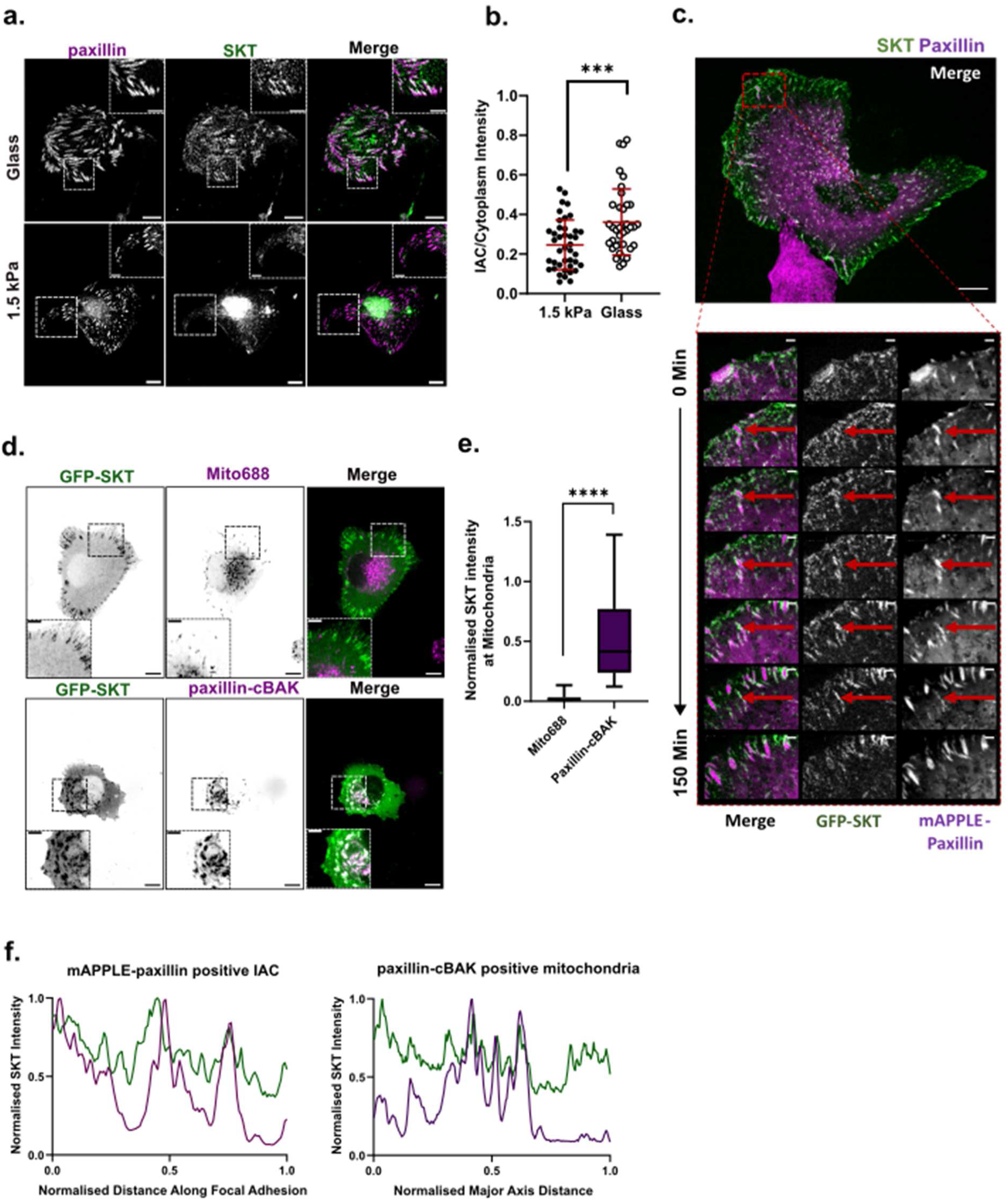
Localisation of SKT at cell-matrix integrin-based adhesions (IAC’s) with paxillin on stiff microenvironments. **a.** Representative images of endogenous localisation of SKT (green) on 1.5 kPa and Glass (≥1 GPa) rigidity substrates, with paxillin (magenta) to visualise IACs in U2OS cells. **b.** Quantification of IAC/Cytoplasm SKT intensity ratios on 1.5 kPa and Glass substrate imaging dishes. **c.** Live cell time-lapse microscopy of U2OS cells expressing GFP-SKT (green) and mAPPLE-paxillin (magenta). Representative images with zoomed insets showing IACs from assembly to disassembly at 0-150 min post-cell spread. **d.** Representative confocal images for the mitochondrial targeting assay showing stable transfected U2OS cells expressing GFP-SKT (green) and cells exposed to a MitoTracker688^TM^ dye (magenta, top panel), or co-transfected with cBAK-paxillin (magenta, bottom panel). **e.** Mean co-localisation of SKT and MitoTracker688^TM^, or cBAK-Paxillin, across the major axis of the total mitochondrial area. **f.** Line scan analysis of SKT and paxillin intensity profiles across the major axis of cBAK-paxillin positive total mitochondrial area, or mAPPLE-paxillin positive lateral IAC length. Results are displayed as mean ± SD and significance was tested using an independent t-test for IAC/Cytoplasm SKT intensity ratios. ***, P ≤ 0.001 (C). Results are displayed as box plots showing the mean, 25^th^, and 75^th^ quartiles ± SD. Significance was tested using an independent t-test corrected with a Mann Whitney test for SKT intensity line scans. ****, P ≤ 0.0001. (F). Scale bar = 10 µm (zoomed insets = 5 µm).

As an adapter protein facilitating key molecular interactions at adhesion sites, paxillin plays an important role in regulating the functions and dynamics of IACs via direct binding and recruitment of a wide variety of IAC-associated proteins including FAK, βPix, and Git1/2^28, 30,31^. To test whether paxillin binds SKT, we used a knock-sideways system in which paxillin was fused to a mitochondrial targeting sequence (cBAK), enabling acute relocalisation of paxillin and associated proteins to mitochondria^32^. Subsequent imaging with the mitochondrial-specific live cell dye, MitoTracker688^TM^, displayed successful targeting of paxillin to the mitochondria (**Fig. S1c**). Following confirmation of the mitochondrial localisation of paxillin, cells were co-transfected with either GFP-SKT and mCherry-paxillin-cBAK, or GFP-SKT and counterstained with MitoTracker688^TM^ (**Fig. 1d**). GFP-SKT was re-localised to the mitochondria with mCherry-paxillin-cBAK and shows significant co-localisation at the mitochondria, whereas the MitoTracker control displays no comparable localisation beyond background signal (**Fig. 1e, f**). In addition, we were able to validate this interaction through co-immunoprecipitation of SKT with GFP-paxillin using GFP-Trap^®^ (**Fig. S1d**). These imaging experiments demonstrate that SKT localises to integrin adhesion complexes in a stiffness-dependent manner and that its recruitment requires an interaction with paxillin.

### SKT is required for adhesion dynamics and cell migration

To examine the role of SKT in regulating the structure and function of IAC’s, we performed immunofluorescence on fixed cells following depletion of SKT expression using two independent siRNAs. Cells were seeded onto fibronectin and collagen coated coverslips and immuno-stained for paxillin. SKT depleted cells displayed a significant decrease in adhesion number in comparison to control cells (siNT) (**Fig. 2a, b**). Qualitative analysis indicated that many paxillin-positive adhesions were localised at the periphery of the cell rather than being spread across the cell’s basal surface as observed in our control cells (**Fig. 2a**). Further analysis of IAC morphology following depletion of SKT revealed an increase in the average size of the remaining adhesions (**Fig. 2a, b**). In addition, these phenotypes were rescued with transient re-expression of GFP-SKT, following siRNA-mediated knockdown of SKT using an siRNA that targets SKT outside of the open reading frame (siSKT_2) (**Fig. 2a, b**).

**Figure 2.**
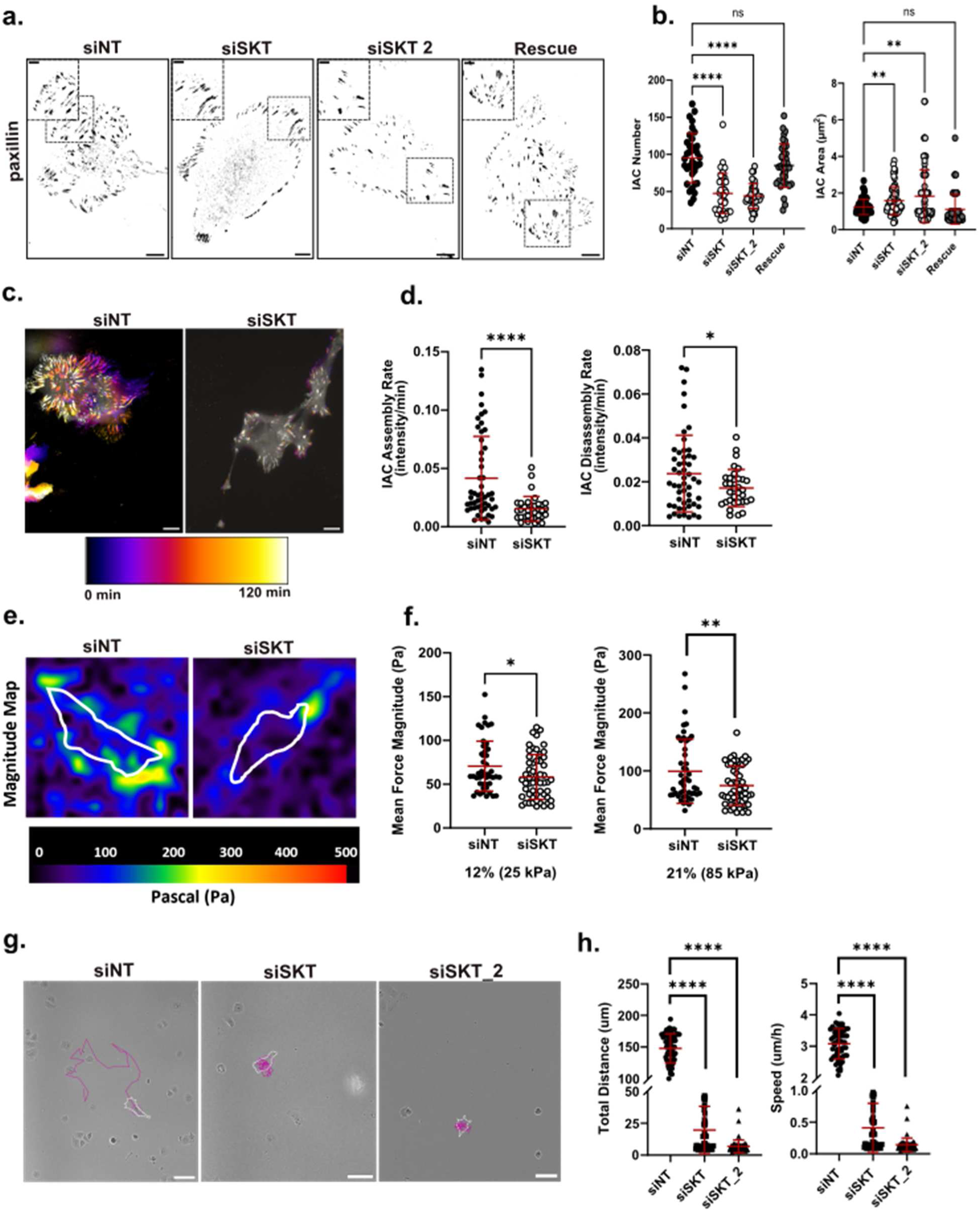
SKT is required for the maintenance of integrin-based adhesion (IAC) function. **a.** Representative images of endogenously localised paxillin using immunofluorescence to visualise IACs on stiff 2D environments in U2OS cells treated with non-targeting control siRNA (siNT), two independent SKT-targeted siRNAs (siSKT and siSKT_2) or rescued with GFP-SKT overexpression (Rescue). **b.** Quantification of FA number and Area in control (siNT), SKT depleted (siSKT and siSKT_2), and GFP-SKT Rescued U2OS cells **c.** Temporal pseudo colour-coded representative images with temporal colour scale for U2OS cells treated with non-targeting control siRNA (siNT) and SKT-targeted siRNA (siSKT), transiently transfected with mAPPLE-paxillin to track IAC dynamics. **d.** Quantification of IAC assembly and disassembly rate in control and SKT depleted cells, where only IACs with both assembly and disassembly rates were included. **e.** Representative Pseudo-coloured image of cell generated traction force magnitude maps on stiff polyacrylamide (PAA) hydrogels using traction force microscopy (TFM) with colour-coded scale bar (Pascals, Pa) included. Cell position outlined in white. **f.** Quantification of mean traction force per cell on stiff PAA hydrogels of 15-30 kPa (12% PAA) and 70-100 kPa (21% PAA). Tractions calculated as averaged individual force vector magnitudes per cell. **g.** Representative bright-field images of single cell random migration in U2OS cells treated with non-targeting control siRNA (siNT), or two independent SKT-targeted siRNAs (siSKT and siSKT_2) with their migration path tracked (purple line) over 48 h. **h.** Quantification of total migration distance and speed in siNT, siSKT, and siSKT_2 treated cells. All results are displayed as mean ± SD. Significance was tested using a one-way Anova for adhesion phenotypes. **, P ≤ 0.01. **** P ≤ 0.0001. (C). Significance was tested using an independent t-test for assembly and disassembly dynamics. *, P ≤ 0.05, ****, P ≤ 0.0001. (E). Significance was tested with independent t-tests for TFM. *, P ≤ 0.05, **, P ≤ 0.01. (G). Significance was tested using a one-way ANOVA for cell migration, ****, P ≤ 0.0001. Scale bar = 10 µm (zoomed insets = 5 µm).

Changes in adhesion size and number are often the product of altered dynamics. To test this the assembly and disassembly of cell-matrix adhesions was examined using time lapse fluorescence microscopy. IACs were less dynamics in SKT knockdown cells, with a significant decrease in both adhesion assembly and disassembly, compared to control cells (**Fig. 2c, d**). Overall, SKT significantly influences the morphological and dynamic characteristics of IAC’s, suggesting a key role in the regulation of cell-matrix adhesion function.

We next asked how changes in adhesion dynamics might also be accompanied by changes in force regulation, ECM sensing, and migration. To probe this, traction force generation was investigated using traction force microscopy on SKT depleted cells cultured on stiff microenvironments. Overall, there was a significant decrease in mean force magnitude exerted when SKT was depleted (**Fig. 2e**). A greater difference in magnitude of force compared to the control was observed as substrate stiffness was increased from 15-30 kPa to 70-100 kPa, resulting in an average difference of 12 Pa and 25 Pa, respectively with SKT depletion (**Fig. 2f**). As traction force and cell migration are intimately linked to substrate rigidity via adhesion dynamics^33, 34^, we sought to examine whether SKT is required for efficient cell migration in stiff microenvironments. Cell tracking analysis of single cell random migration displayed a significant reduction in both the migration distance and speed when SKT was depleted (**Fig. 2g, h**). These findings establish SKT as a critical regulator of adhesion turnover, enabling efficient IAC assembly and disassembly required for force generation and directed cell migration.

### SKT activates mTORC2 signalling at cell-matrix adhesions

To investigate the molecular mechanisms underlying SKT function, we identified its binding partners using GFP-Trap® immunoprecipitation followed by mass spectrometry (**Table S1**). Proteins were classified as SKT interactors based on unique peptide signatures enriched relative to a GFP-only control. Serval adhesion- and cytoskeletal-associated proteins were identified, such as Tensin1, GIT1, Liprin-α1 and filamin A^35–37^ (**Fig. 3a**). Beyond canonical adhesion-associated interactors, SKT co-precipitated with core components of mTORC2 (mTOR, RICTOR, mSIN1), with no detectable mTORC1 subunits, indicating a selective association (**Fig 3a, Table S1**).

**Figure 3.**
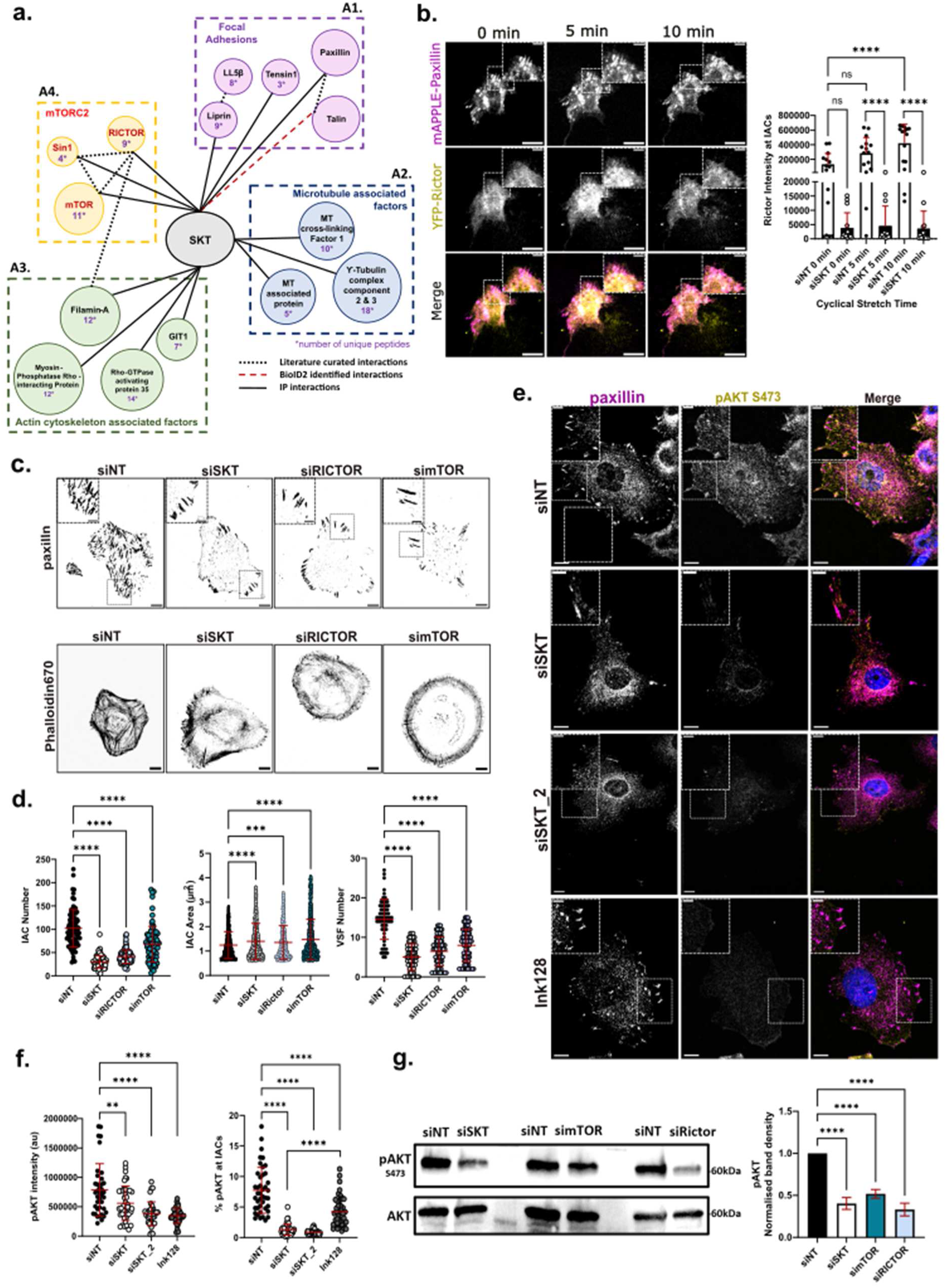
SKT is a novel interactor of mTORC2 at cell-matrix adhesions. **a.** Interaction map of SKT interactors based on GFT-Trap immunoprecipitation (IP) and mass spectrometry. **A1.** Identified proteins associated with Integrin-based adhesions (focal adhesions). **A2.** Identified proteins associated with microtubules and the centrosome. **A3.** Identified proteins associated with the actin cytoskeleton. **A4.** Identified proteins associated with the mechanistic target of rapamycin complex 2 (mTORC2). **b.** Representative images of non-targeted siRNA (siNT) treated U2OS cells under uniaxial stretching with quantification of YFP-Rictor intensity at 0-, 5-, and 10-min post stretching. **c.** Representative images of endogenous paxillin (top panel) to visualise adhesions and Acti-stain^TM^ Phalloidin-670 to visualise the actin cytoskeleton (bottom panel) in U2OS cells treated with siNT, siSKT, mTOR-targeted siRNA (simTOR), and RICTOR-targeted siRNA (siRICTOR). **d.** Quantification of integrin-based adhesion number and area, and ventral stress fibre number in siNT, siSKT, simTOR, and siRICTOR treated cells. **e.** Representative images of endogenous pAkt in siNT, siSKT, siSKT_2, and dual mTORC1/2 inhibitor, INK-128, treated U2OS cells. **f.** Quantification of whole cell pAkt intensity in siNT, siSKT, siSKT_2, and INK-128 treated cells, and pAkt localised at IACs in siNT, siSKT, siSKT_2, and Ink-128 treated cells. pAkt intensities at IACs were normalised to whole cell pAkt intensity to give percentage pAkt at IAC sites. **g.** Representative western blot and quantification of pAkt band intensity of siNT, siSKT, simTOR, and siRICTOR treated U2OS cells. Blots probed for pAkt (S473), and Total Akt as a control. Results are displayed as mean ± SD per cell and significance was tested using a one-way ANOVA for comparisons of YFP-Rictor intensity. ****, P ≤ 0.0001, ns, P ≥ 0.05. Significance was tested with a two-way ANOVA for adhesion and actin quantifications. ****, P ≤ 0.0001, ***, P ≤ 0.001. Significance was tested with one-way ANOVA, corrected with a Kruskal-Wallis test where data is not normally distributed for pAKT quantification, **, P ≤ 0.01, ****, P ≤ 0.0001. Significance was tested with a one-way ANOVA for pAKT band intensity of western blots. ****, P ≤ 0.0001. Scale bar = 10 µm (zoomed insets = 5 µm).

To validate the co-localisation of SKT and mTORC2 and determine whether this is directly associated with SKT’s function at cell-matrix adhesions, we explored the localisation and recruitment of mTORC2, focussing on the recruitment of RICTOR to IACs. Since our previous results suggest SKT recruitment to IACs is mechanosensitive, we examined RICTOR recruitment to IACs in the presence and absence of SKT in cells exposed to cyclical uniaxial stretching, which is known to affect IACs^38^. We observed an increase in the recruitment of RICTOR to IACs after 10 minutes of cyclic stretching in the control cells, which was not observed in the SKT knockdown cells (**Fig. 3b; Fig. S2a**).

To explore whether the phenotypes observed after SKT depletion are mediated *via* mTORC2/RICTOR, we depleted either mTOR or RICTOR using siRNA (simTOR and siRICTOR) and examined IACs in these cells. Depletion of either resulted in a similar phenotype to siSKT-transfected cells: a reduction in IAC number and an increase in the size of the remaining adhesions when all three proteins were independently depleted (**Fig. 3c, d**). We also observed distinct differences in cytoskeletal organisation in all knockdown conditions, with a significant reduction in ventral stress fibres compared to control cells (**Fig. 3c, d**). Together, these data suggest a shared mechanism between SKT and mTORC2 in regulating IAC and cytoskeletal dynamics in response to mechanical cues.

One functional impact of mTORC2 is the direct phosphorylation of Akt at serine 473 to promote downstream signalling^39–41^. To determine whether SKT influences mTORC2 function, we quantified phospho-Akt (Ser473) as a functional marker of mTORC2 activity. Both immunofluorescence examining phospho-Akt localisation and Western Blot analysis to measure phospho-Akt protein levels indicated a significant reduction in the percentage of phospho-Akt present at IACs and whole cell phospho-Akt, respectively following SKT depletion (**Fig. 3e-g**). These results were comparable to the effect on phospho-Akt present at IACs following treatment with the dual mTORC1/2 ATP competitive inhibitor, Ink128 (**Fig. 3e, f; Fig. S2b**), and phospho-Akt protein levels following siRNA-mediated silencing of mTORC2 (**Fig. 3g**). Together, these data suggest that SKT recruits mTORC2 to IACs and that SKT is required for downstream signalling/function of mTORC2.

### Adhesion Signalling is directly linked to Glucose Metabolism levels

As mTORC2 can regulate glucose uptake and glycolysis through its activation of Akt and subsequent downstream signalling^42–46^, we reasoned that SKT may play a significant role in the regulation of glucose metabolism to influence metabolic flux in response to matrix mechanics. First, to identify a more relevant model for investigating the physiological function of SKT, we utilised the GEPIA webserver using data from The Cancer Genome Atlas (TCGA) and Genotype-Tissue expression (GTEx) databases to investigate SKT expression in various malignancies. SKT is expressed at higher levels in tumour vs normal tissue and was found to be associated with a poorer prognosis in Pancreatic Ductal Adenocarcinoma (PDAC) (**Fig. 4a**). As desmoplasia is a prominent feature of the PDAC microenvironment (REF), we decided that further investigation into SKT’s function in response a stiffer matrix is more clinically relevant using a PDAC cell line.

**Figure 4.**
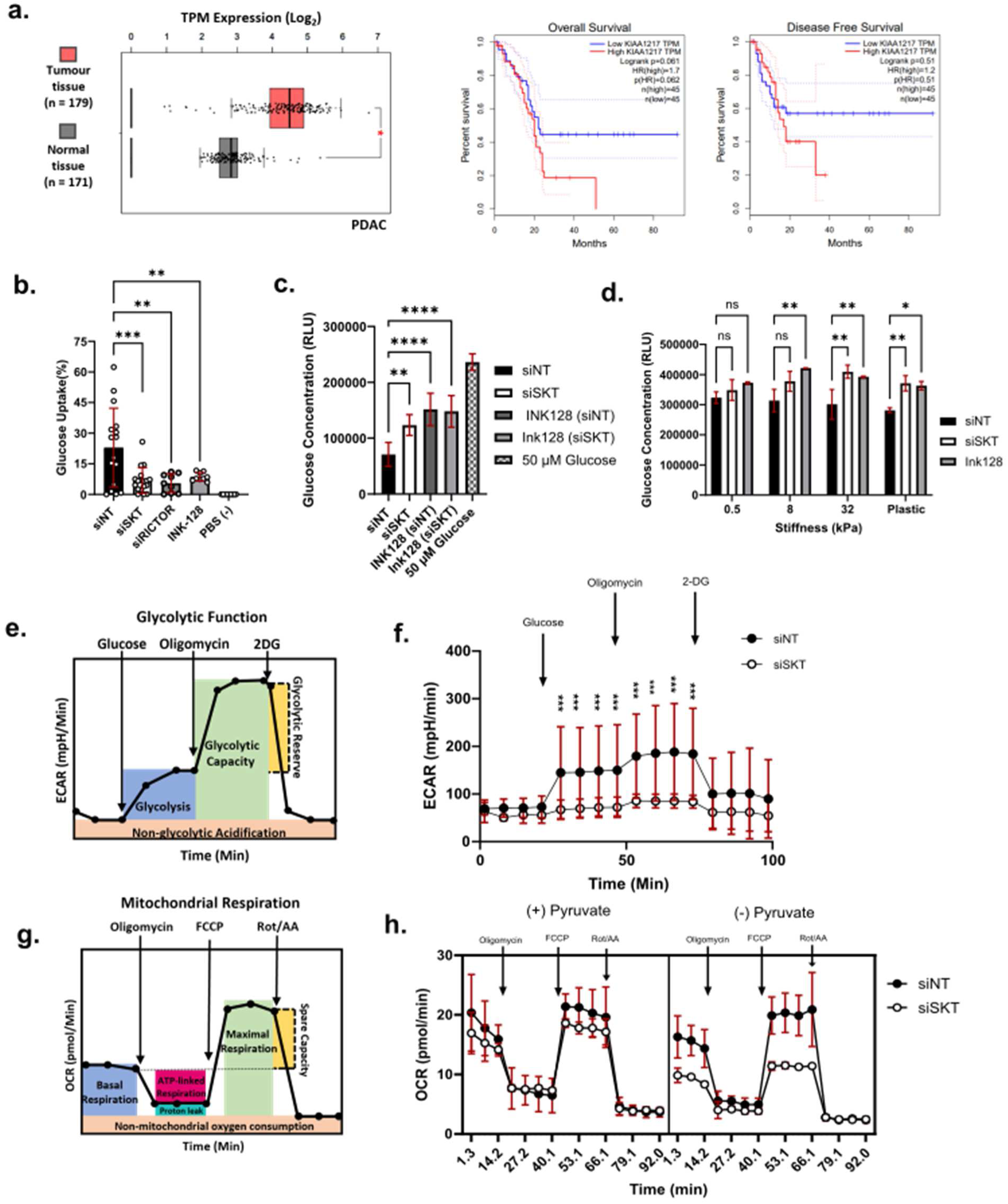
Depletion of SKT significantly influences glucose metabolism. **a.** Transcript per million (TPM) expression of SKT in tumour (PDAC) tissue compared to healthy tissue counterparts. Data was obtained from patients with PDAC using the GEPIA webserver from RNA sequencing data obtained through two cancer databases, the cancer genome atlas (TCGA) and the genotype-tissue expression database (GTEx). A Log_2_ expression cut off = 1.5 and p-value cut off = 0.01 were used to filter the samples matching TCGA normal and GTEx data. Kaplan-Meier analysis showing the correlation between SKT (*KIAA1217*) expression and PDAC prognosis. Both overall survival and disease-free survival were analysed using the GEPIA webserver (http://gepia.cancer-pku.cn/), and p-value cut off = 0.01 was applied.**b.** Quantification of glucose uptake (represented by uptake of 2-deoxyglucose, 2DG) in Panc-1 cells treated with control, non-targeting siRNA (siNT), SKT-targeted siRNA (siSKT), RICTOR-targeted siRNA (siRICTOR), and dual mTORC1/2 inhibitor Ink-128 using the Glucose Uptake^TM^ assay (Promega). PBS was used as a negative control. Uptake percentages were calculated relative to the highest glucose uptake value observed in Panc-1 cells following 30 minutes of 1 mM 2DG incubation. **c.** Quantification of glucose consumption using the Glucose-Glo^TM^ assay (Promega) in Panc-1 cells treated with siNT and siSKT in the absence of INK-128, and siNT and siSKT in combination with INK-128. Data shown as relative light units (RLU) from absorbance measurements following 48 h incubation of cells with 5 mM glucose. A total glucose concentration of 5 mM was serially diluted to 50 µM to calulate relative glucose concentrations **d.** Quantification of glucose consumption using the Glucose-Glo^TM^ assay (Promega) in Panc-1 cells treated with siNT, siSKT, and treated with INK-128 on 0.5-, 8-, and 32-kPa substrates. **e.** Schematic diagram of Seahorse XFe24 metabolic analyser glycolytic stress test read out in response to sequential injections of 10 mM glucose, 1 µM Oligomycin, and 50 mM 2-deoxyglucose (2DG). **f.** Quantification of extracellular acidification rate (ECAR) and the cellular response to glucose, Oligomycin, and 2DG in Panc-1 cells treated with siNT and siSKT. **g.** Schematic diagram of Seahorse XFe24 metabolic analyser mitochondrial stress test read out in response to sequential injections of 1 µM Oligomycin, 2 µM Carbonyl cyanide-4 (trifluoromethoxy) phenylhydrazone (FCCP), and 0.5 µM Rotenone and Antimycin A (Rot/AA). **h**. Quantification of oxygen consumption rate (OCR) and the cellular response to Oligomycin, FCCP, and Rot/AA in Panc-1 cells treated with siNT and siSKT with or without the addition of pyruvate supplemented in the media. All results are displayed as mean ± SD per cell. Significance was tested with a one-way ANOVA for glucose uptake. **, P ≤ 0.01, ***, P ≤ 0.001. Significance was tested with a one-way ANOVA for glucose consumption between siNT and siSKT. **, P = 0.0054, ****, P ≤ 0.0001. Significance was tested with a two-way ANOVA for glucose consumption comparisons between stiffness substrates. *, P ≤ 0.05, **, P ≤ 0.01. Significance was tested with a two-way ANOVA for the glycolytic and mitochondrial stress test. ***, P ≤ 0.001, **** P ≤ 0.0001, ns, P ≥ 0.05.

We then assessed glucose uptake using the Glucose Uptake-Glo^TM^ assay (Promega). This assay uses a nonradioactive, homogenous bioluminescent method based on the detection of 2-deoxyglucose-6-phosphate (2DG6P), via uptake of 2-deoxyglucose (2DG) (**Fig. S2e**). In response to SKT and RICTOR depletion, 2DG uptake was significantly reduced by approximately 70% and 75%, respectively, relative to the control. Similarly, we observed a 62% decrease when mTOR signalling was inhibited with the dual mTORC1/2 ATP competitive inhibitor Ink-128 (**Fig.4b**). To validate our findings, we analysed glucose consumption to detect cellular glucose levels (**Fig. S2f**) following depletion of SKT, and inhibition of mTOR signalling (with Ink-128). In both conditions, we also observed a significant reduction in glucose consumption when compared to the control (**Fig. 4c**).

Our previous results show SKT and mTORC2 recruitment to IACs is mechanosensitive. We next examined whether their recruitment in response to external strain altered glucose metabolism by measuring glucose consumption in cells on substrates of different stiffness. As substrate stiffness was increased, control cells displayed a trend toward greater glucose consumption (indicated by a decrease in glucose concentration) (**Fig. 4d**). Following depletion of SKT or Inhibition of mTOR, a higher concentration of glucose remains unconsumed on substrates resembling a stiffer-desmoplastic-tumor matrix (8-32kPa) and plastic, when compared to the control. (**Fig. 4d**). These results revealed a functional importance of mTORC2 and SKT in mechanically regulated glucose consumption in stiffer microenvironments.

Elevated glucose uptake is a crucial hallmark of increased glycolysis in cancer cells^47^, therefore, to elucidate the function of SKT in glucose metabolism and metabolic flux, we next sought to examine SKT’s role in glycolysis and/or oxidative phosphorylation (OxPhos). To assess glycolytic activity, we performed a Seahorse glycolytic stress test to measure extracellular acidification rate (ECAR). Sequential addition of glucose, oligomycin, and 2-deoxyglucose enabled quantification of basal glycolysis, glycolytic capacity, and non-glycolytic acidification (**Fig. 4e**). When SKT was depleted the mean ECAR did not increase beyond 90 mpH/min, compared to 190 mpH/min in control cells, a significant reduction in the glycolytic capacity of SKT-depleted cells. Compared to the control, SKT depleted cells displayed a decrease in ECAR at each stage of the analysis (**Fig. 4f**) indicating a significant decrease in glycolysis, glycolytic capacity, and glycolytic reserve upon SKT depletion.

To determine whether the metabolic effects of SKT depletion extend beyond glycolysis, we assessed mitochondrial respiration using a Seahorse mitochondrial stress test (**Fig. 4g**). In the presence of exogenous pyruvate, which uncouples oxidative phosphorylation from glycolytic input, SKT-depleted cells displayed oxygen consumption rates comparable to controls across all respiratory phases (oligomycin, FCCP, and Rot/AA treatments; **Fig. 4h**). These findings indicate that loss of SKT does not impair mitochondrial respiratory capacity and that SKT–mTORC2 signalling primarily affects glycolytic, rather than oxidative metabolism^48–51^. To provide a holistic view of metabolic flux in relation to the role of SKT in glycolysis, we repeated the mitochondrial stress test but in the absence of pyruvate. Here, we observed a greater decrease in basal and maximal respiration (**Fig. 4h**). These findings support a glycolysis-specific role for SKT–mTORC2 signalling.

### Tumour invasion and growth is significantly affected by SKT expression

Metabolic rate adaptation and adhesion dependent invasion are key hallmarks of tumour progression^52^. This prompted us to test if there was a direct effect of SKT expression level on tumour formation and growth using the chick embryo chorioallantoic membrane (CAM) model. We generated stably expressing GFP-SKT cells to simulate high SKT expression and used siRNA knockdown to reduce expression. Overexpression of GFP-SKT resulted in higher tumour formation on CAMs than wild type (**Fig. 5a, b**) whereas reduction of SKT expression level had the opposite effect and showed significantly reduced tumour number and mass compared to the control (**Fig. 5c, d**). Collectively, these data identify SKT as a mechano-metabolic regulator whose elevated expression promotes tumour growth *in vivo*.

**Figure 5.**
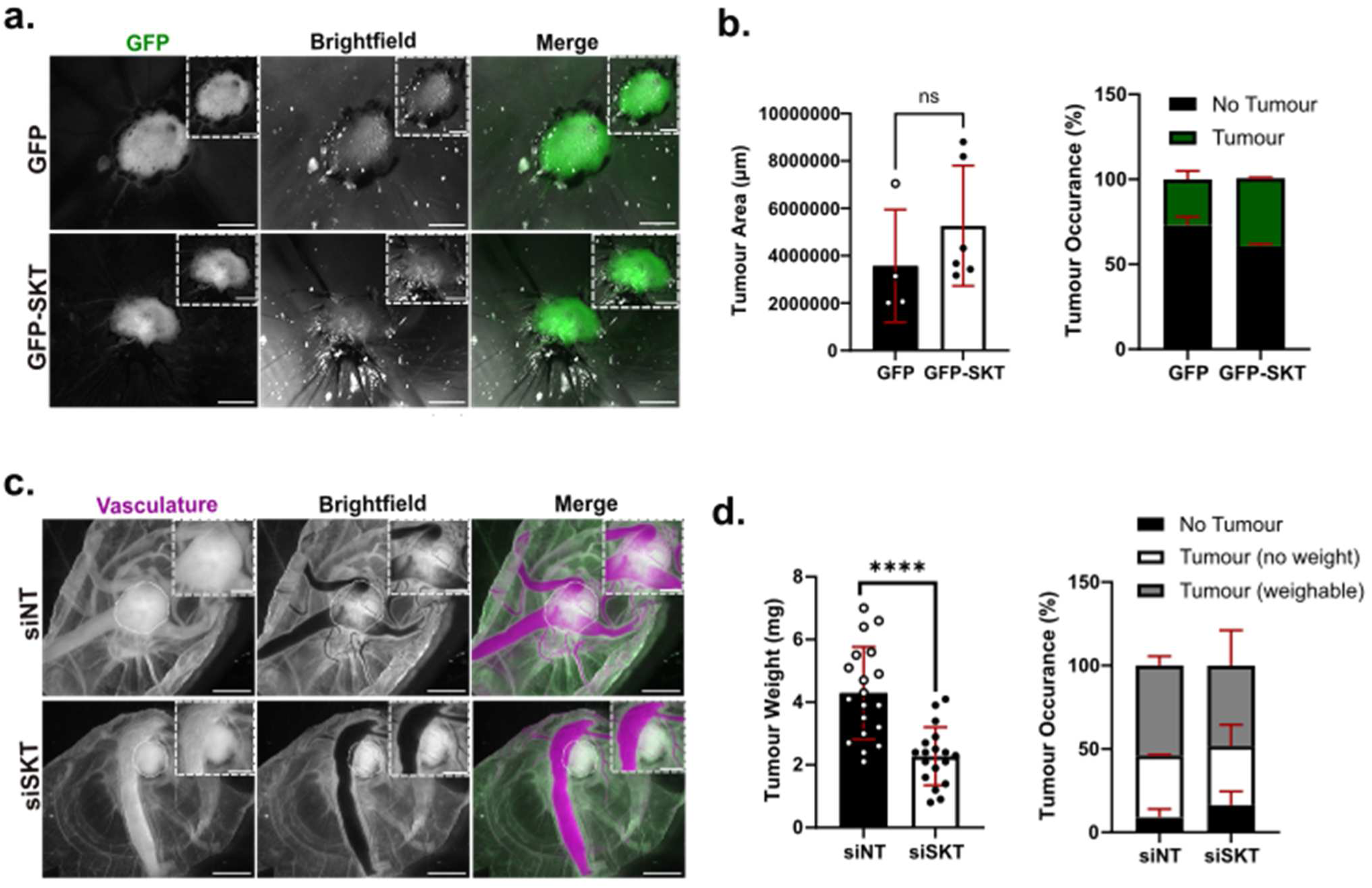
Expression of SKT influences tumour progression and maintenance. **a.** Representative in vivo imaging of GFP control and GFP-SKT Panc-1 tumours formed within the chorioallantoic membrane (CAM) of fertilised chick embryos. **b.** Quantification of tumour area (□m^2^) and percentage tumour occurrence from GFP and GFP-SKT expressing Panc-1 cells in CAM. **c.** Representative in vivo imaging of non-targeted siRNA (siNT) and SKT-targeted siRNA (siSKT) treated Panc-1 tumours within the CAM of fertilised chick embryos. **d.** Quantification of tumour weight (mg) and percentage tumour occurrence from Panc-1 cells in CAM. All results are displayed as mean ± SD per tumour formed. Significance was tested using an independent t-test for tumour weight and area. ****, P ≤ 0.0001. Scale Bar = 100 µm (zoomed insets = 50 □m).

## Discussion

The tumour microenvironment (TME) is populated with many different cell types including tumour cells and cancer-associated fibroblast which contribute to the common conversion of the physiological matrix to a stiffer and denser desmoplastic tumour matrix. Desmoplasia is characterised by an increase in α-smooth muscle actin positive fibroblasts, that produce an abundance of ECM proteins including types I and V collagen and fibronectin^25, 26^. This significantly increases ECM stiffness, leading to changes in IACs and increased ATP-dependent actomyosin contractility^27, 28^. This process reinforces many aggressive characteristics observed across tumour types and has been implicated in chemotherapeutic resistance and metabolic plasticity^29, 53–55^.

Here we show that the previously uncharacterised protein SKT is a novel cell-matrix adhesion protein preferentially enriched on stiffer matrices. We identify a role for SKT in the regulation of IAC dynamics and demonstrate that SKT recruits mTORC2, a pivotal signalling complex in cellular metabolism, growth and survival, to IACs in response to extrinsic force. Loss of mTORC2/RICTOR mirrors the observed changes to IACs without SKT and together they are required for activation of Akt and regulation of glucose metabolism within our model.

### SKT/mTORC2 recruitment and role at adhesions

We show that SKT is a novel paxillin associated protein. As an adapter protein facilitating key molecular interactions at adhesion sites, paxillin regulates the functions and dynamics of IACs via direct binding of proteins such as vinculin, FAK, Git and talin through its leucine rich LD motifs^28, 30, 31, 56^.

Paxillin function is at least in part regulated by phosphorylation and ubiquitination, at multiple sites throughout the protein^31, 56^, and disruption to paxillin phosphorylation has been demonstrated to significantly disrupt IAC formation and disassembly^30, 57, 58^.

Interestingly a recent phosphoproteomics study in neuroblastoma cells identified paxillin Ser85 and S322 as potentially phosphorylated by mTORC2^59^. Paxillin Ser85 phosphorylation has been shown to be dependent on cell adhesion and required for adhesion formation and cell migration^60^. Future studies will have to address the mTOR phosphorylation targets in IACs to elucidate further details of its role in adhesion dynamics.

### mTORC2 signalling at adhesions

In our study, SKT mediated recruitment of mTORC2 to adhesion sites is required for activation of its downstream target Akt. Activation of mTORC2 has been demonstrated in response to growth factors/hormones^61^ and membrane tension^61–63^, but no direct link to ECM sensing has yet been made. A previous study showed that mTORC2 is recruited to the plasma membrane, to bring it into proximity with its downstream effectors and their additional activators, such as Akt and PI3K/PDK1^64^. We can show increased mTORC2 localisation to IAC sites when cells are subjected to mechanical force or seeded onto stiff substrates, suggesting that ECM signals sensed at IACs are integrated and converted into signals that regulate metabolic demand of the cell. SKT could serve as a rheostat that controls metabolic adaptations of cells in response to changes in their matrix environment.

Akt is a Ser/Thr protein kinase that has been extensively studied for its role in regulating glucose metabolism following activation by mTORC2^64–66^. Increased matrix stiffness has been shown to increase energy demand and glucose consumption^67, 68^. Park et al., (2020) report an increase in glycolysis when stiffness is increased. They demonstrate a higher expression of the glycolytic enzymes hexokinase-2 and phosphofructokinase in human bronchial epithelial cells when they are plated on stiff (collagen-coated glass) substrates in comparison to soft (Pa range elastic collagen) substrates, and link this to the mechanical regulation of stress fibres^19^. This is also supported by Ponce et al., (2021), who demonstrate an increase in lactate production and glucose uptake via GLUT1 in mammary fibroblasts when stiffness is increased from 0.2 kPa to 32 kPa^21^. Through its activation of Akt, mTORC2 is a key regulator of both hexokinase-2 and phosphofructokinase expression^19, 20, 43, 44, 69^ and can significantly influence glucose metabolism via regulation of glucose transporters like GLUT1^43, 44^.

These previous findings support our hypothesis that recruitment of SKT and mTORC2 (to IACs) plays a vital role in glucose metabolism when stiffness is increased. Until this study, the mechanisms that linked mTORC2 regulated glucose metabolism (glucose uptake and glycolysis) with ECM stiffness were poorly understood, and exactly how the cells sensed changes in ECM stiffness and translated this to a glycolytic response were unknown.

### Tumour growth

Key cellular metabolic pathways, including glucose metabolism, are significantly dysregulated in cancer and provide a central mechanism for the upregulation of cellular energetics, required for tumour proliferation and tumorigenesis^21, 70^. ECM stiffness significantly influences the malignancy of tumour cells and has been shown to be able to reprogram their metabolic profiles^67, 68^. For example, increased ECM stiffness can significantly increase glycolysis and influence glycolytic flux (glycolysis, glycolytic capacity, and glycolytic reserve)^67, 71^. A high glycolytic capacity provides rapid ATP production to facilitate the energetic demands of cancer cells, including their reorganisation of the actin cytoskeleton and maintenance of force generation, and provides cells with a growth advantage over their neighbours^72^.

Dysregulation of cell adhesion to the ECM has been shown to strongly influence glycolytic flux although via an unknown mechanism^23^. We propose that SKT plays a pivotal role in this response. Here, we show that depletion of SKT and mTORC2 significantly reduces glycolysis in Panc-1 cells and begin to explore the implications of this on tumour growth. Epstein et al., (2014) highlight the role of glycolytic flux in the cells ability to respond to fluctuating energetic demands and demonstrate the requirement for high cellular glycolytic reserves^73, 74^. Our data shows that SKT depleted cells have a lower glycolytic capacity and are therefore less capable of responding to a rapid demand in energy as their glycolytic function is already at the cell’s theoretical maximum. This suggests that without SKT, cells lack the ability to metabolically respond to their environment and lose their glycolytic advantage for survival within the tumour microenvironment. To test this, we used the CAM model and show that when SKT was overexpressed, tumour growth was increased in comparison to the control, and the opposite was found when SKT was depleted. Together, this demonstrates a novel function for SKT in regulating metabolic plasticity and confirms a role for SKT in tumour maintenance and progression. The novel functions identified for SKT in IACs, mTORC2 signalling, and glycolysis unveils a promising avenue for therapeutic exploration in cancer treatment and highlights a novel signalling axis between ECM stiffness and cancer metabolism.

## Methods

### Cell Culture

U2OS and PANC-1 cell lines were obtained from ATCC. Cells were cultured in a humidified Incubator at 37°C with 5% CO2 in DMEM GlutaMax^TM^ media (Thermo Fisher Scientific, Cat #31966021) containing 4.5 g/L D-Glucose and Sodium Pyruvate and supplemented with 10% Foetal Bovine Serum (Thermo Fisher Scientific, Cat #A4766801) and 1% Penicillin and Streptomycin (Sigma-Aldrich, Cat #15140122).

### Subculture

Cell Culture media was aspirated, and cells were washed with sterile PBS. 2.5mL of 0.05% Trypsin-EDTA solution (Thermo Fisher Scientific, Cat #15400054) per 75cm^2^ area was added, and the cells were incubated in a humidified Incubator at 37°C with 5% CO2 for 5min untill >70% of cells were seen to have detached under bright field observation. Complete cell culture media was then added to the cell suspension at 5-fold volume and agitated to ensure a homogenous cell suspension solution. An appropriate volume was then taken and seeded into a fresh cell culture vessel and incubated at 37°C with 5% CO2.

### siRNA Transfection

For transfection of 4 x 10^5^ cells seeded into a 6-well plate; 30 nM siRNA and INTERFERin^®^ transfection reagent (Polyplus, Cat #101000028) were combined with 200 μL of Opti-MEM^®^ (reduced serum media, Cat #11058021). The siRNA/transfection reagent mix was then incubated at room temperature for 10 minutes. During incubation, cell culture media was aspirated from the cells and 1.8mL was re-added. Following incubation, the transfection solution was added to the cells dropwise. For all knockdowns performed, cells were subject to a siRNA treatment at 24 and 72 hours post seeding. The cells were then evaluated in each assay between 72-96 hours following the first treatment.

### DNA Transfection

Depending on the desired assay scale, cells were grown to ∼70% confluence in a 6-well plate using standard cell culture conditions (as previously detailed). 1 μg of plasmid DNA and 4 μL jetPRIME^®^ transfection reagent (Polyplus, Cat #101000046) were combined with 100 μL jetPRIME^®^ Buffer (Polyplus, sterile filtered 0.2μm, Cat #201000003) and mixed. Transfection was then carried out in accordance with the manufacturer’s instructions.

### Drug Treatments

#### Sapanisertib

For Sapanisertib (Ink-128, MedChem Express. Cat #1224844-38-5) treatment, cells were cultured to 70% confluency in 6-well plates. 2.5 mM stock solution was then serial diluted in 2 mL DMEM to 20 μM, and again to 200nM in 2 ml 10% FBS supplemented DMEM GlutaMax^TM^ cell culture medium. Cells were treated with Ink-128 for 24-48 h prior to experimental analysis. If re-plated, cells were detached using 1 x Trypsin-EDTA (0.05%) and allowed to attach to fibronectin (10 µg/mL) and collagen (50 µg/mL) coated coverslips. 200 nM Ink-128 containing DMEM was then reapplied to cells prior to further incubation and fixation to maintain sustained treatment for the appropriate time.

### Immunofluorescence

13mm coverslips were washed three times in 70% ethanol and then dried between two sheets of whatman filter paper in a laminar flow hood. The coverslips were then sterilised via autoclaving and dried in a drying cabinet, prior to being stored in a clean, sealed container until use. All coverslips used were then coated with fibronectin (10 µg/mL) and collagen (50 µg/mL) at 4°C for a minimum of 4 h.

Cell culture media was aspirated from cells and coverslip, and then the coverslips were incubated in 4% paraformaldehyde (PFA, w/v) (Sigma-Aldrich, Cat #158127500G) solution in PBS for 15 minutes. Following removal of the PFA, the samples were washed three times in PBS and permeabilised via treatment with 0.3% Triton X-100 solution (v/v) (Sigma-Aldrich, Cat #T9284) in PBS for a further 15 minutes. The samples were then washed three times with PBS and blocked with 10% bovine serum albumin (BSA, cohn fraction V, w/v) (First Link UK Ltd, Cat #4100450) for 30 minutes. The blocking buffer was then aspirated off, and the primary antibodies were diluted in 1% BSA in PBS, before being incubated on the sample for 1 hour. The sample was washed three times in PBS, and the secondary antibodies were diluted in 1% BSA in PBS. The sample was then incubated with secondary antibody for 1 hour under foil. All incubation steps were carried out at room temperature. If a Phalloidin stain and/or DAPI staining were required, they were also incubated on the sample at this stage (the final 10 minutes for the incubation of DAPI). The sample was then washed three times in PBS and kept in PBS untill mounting.

Excess PBS was removed from samples, and the coverslips were dabbed on white tissue paper to remove and residual PBS. The coverslips were then mounted onto glass slides using ProLong^TM^ Gold antifade mounting agent (Thermo Fisher Scientific, Cat #P36984).

### Fluorescent Microscopy

#### Spinning Disk Confocal Microscopy

All live and fixed cell imaging was captured using a Marianas 3i spinning disk confocal microscope (Intelligent Imaging Innovations, Inc). Images were captured with a sCMOS camera (ORCA-Fusion, Hamamatsu). For each corresponding fluorophore, the appropriate laser configuration and line chosen from 405nm 488nm, 561nm, and 647nm was selected using a suitable emission band pass filter (Table 2.7). All images were captured using a αPlan-Apochromat 63x/1.4NA oil objective lens (Zeiss). For all fixed cell immunofluorescence, z-stack images were captured. For live cell, single z-plane images were captured, and the temperature was set at 37° with 5% CO_2_.

#### Time-lapse microscopy

For focal adhesion turnover and localisation experiments imaging was captured using a Marianas 3i spinning disk confocal microscope (Intelligent Imaging Innovations, Inc), and images were captured with a CMOSS camera (ORCA-Fusion, Hamamatsu). Single Z-stack images were captured using a αPlan-Apochromat 63x/1.4NA oil objective lens (Zeiss). For imaging of mAPPLE-paxillin and GFP-SKT two channels were captured: one channel detecting 495-555nm range under a 488nm excitation with a 300ms exposure, and a second channel detecting a 544-690nm range under 561nm excitation with a 300ms exposure. For focal adhesion turnover, mAPPLE-paxilin time-lapse images were captured every 3 minutes for a duration of 180 minutes (3 hours) to obverse focal adhesion assembly and disassembly events. For cell spreading and GFP-SKT localisation, mAPPLE-paxillin and GFP-SKT time-lapse movies were captured every 5 minutes for a duration of 4 hours.

### Traction Force Microscopy

250 µL of 0.1M NaOH solution was added to the centre of a 13mm glass bottomed imaging dish (MatTek) and incubated for 5 minutes at room temperature. The NaOH was removed and 200 µL of (3-Aminopropyl)-trimethoxysilane (APES) was added for 2.5 minutes. 2 mL of PBS was then immediately added to dilute the APES and stop the reaction, the APES/PBS mixture was then removed. The dishes were then washed twice in MilliQ water prior to adding 400 µL of 0.5% glutaraldehyde solution and incubated for 30 minutes. The dishes were then re-washed twice with MilliQ water, and a 70% ethanol bath for 1 hour. The dishes were then air dried under a tissue culture laminar flow hood prior to use.

Protogel (30% solution at 37:1 ratio of acrylamide:bis-acrylamide) (Sigma-Aldrich, Cat #SLCQ1813) was diluted in PBS to a concentration of 3% (v/v), 12% (v/v), and 21% (v/v). Fluorescent beads with an excitation wavelength of 488nm (Molecular Probes, Cat #F8811) were then mixed into the protogel solution at a ratio of 1:25 by vortex for 2 minutes. The solution was then de=gassed at 400-500mbar for 10 minutes. 10% ammonium persulfate (APS) and tetramethylethylenediamine (TEMED) was added and mixed via gentle pipetting, with care taken to prevent the introduction of air bubbles. 8 µL of the solution was then added to each of the pre-prepared MatTek imaging dishes and covered with a clean 13mm glass coverslip. The dish was then turned up-side down and allowed to set for 30 minutes. The glass coverslip was then removed using forceps and the gel was washed twice in PBS and either used immediately or stored at 4°C immersed in PBS until use.

A 0.2mg/ml silfo-SANPAH solution was freshly prepared, and 200 µL was added to cover the PAA hydrogels. The gels were then exposed to a 365nm UV light source at 10 cm for 10 minutes, before being rinsed three times with 50 mM HEPES solution (pH 8.5). 10 µg/mL fibronectin and 50 µg/mL collagen diluted in the same HEPES solution was added to the functionalised gels and incubated overnight at 4°C. The following day, the gels were washed twice in PBS and sterilised by UV exposure for 20 minutes in a tissue culture laminar flow hood.

1 x 10^3^ U2OS cells were seeded onto the pre-coated PAA gels 24 hours prior to imaging and immediately after gel sterilisation. The stressed state (cells attached) of the gels was imaged using a Marianas 3i spinning disk confocal microscope (Intelligent Imaging Innovations, Inc) and captured using a CMOS camera (ORCA-Fusion, Hamamatsu). Cells were identified by brightfield illumination, and a z-stack encompassing the gels (as indicated by the fluorescent beads) captured under 488 laser illumination. Following the initial acquisition, the cells XYZ positions were saved, and the dishes were treated with 20% SDS to detach the cells from the gel. The gels were then re-imaged at the corresponding XYZ positions. All images were captured using a αPlan-Apochromat 63x/1.4NA oil objective lens (Zeiss) with the lowest possible laser power at 1%.

### Paxillin knock-sideways

Knock-sideways constructs were generated by the Ballestrem lab as described by Atherton et al., (2020)^32^. In brief, PCR was used to generate the 108-bp mitochondrial targeting sequence from cBAK, and site directed mutagenesis was used to remove the stop codon of paxillin and introduce a complementary restriction site to the cBAK fragment. The cBAK fragment was then ligated onto paxillin and cloned into an mCherry (Clontech) C1 vector by restriction digest. To validate the re-localisation of paxillin to the mitochondrial membrane, cells were transiently transfected with mCherry-paxillin-cBAK 24 prior to imaging. Cells were then exposed to 20 nM BioTracker 488 green mitochondrial dye (Sigma-Aldrich, Cat #SCT136, in DMSO) for 20 minutes at 37°C before imaging. Cells were then washed in DMEM GlutaMax^TM^ media (Thermo Fisher Scientific, Cat #31966021) supplemented with 10% FBS. For analysis of SKT localisation, U2OS cells stably expressing GFP-SKT were transiently transfected with mCherry-paxillin-cBAK and incubated for 5 hours before the cell culture medium was replaced. To ensure SKT does not initially localise to the mitochondria, GFP-SKT expressing U2OS cells were exposed to 20 nM BioTracker 633 red mitochondrial dye (Sigma-Aldrich, Cat #SCT137, in DMSO) for 20 minutes at 37°C immediately prior to imaging. All Imaging was then conducted using a Marianas 3i spinning disk confocal microscope (Intelligent Imaging Innovations, Inc). Images were captured using a CMOSS camera (ORCA-Fusion, Hamamatsu), with two channels: one channel detecting 495-555nm range under a 488nm excitation with a 150ms exposure, and a second channel detecting a 544-690nm range under 561nm excitation with a 150ms exposure. Colocalisation analysis was then conducted using FIJI (ImageJ) and intensity line scan profiles were produced.

### Cell Stretching

Cells were transiently transfected with mAPPLE-paxillin and YFP-RICTOR as previously stated and polydimethylsiloxane (PDMS) cell stretching membranes were coated with 10 μg/mL Fibronectin and 50 µg/mL collagen (in PBS) and incubated at 4°C. 6 hours post transfection, stretching membranes were washed twice in PBS and cells were reseeded onto the PDMS cell stretching membranes and incubated at 37°C for approximately 16 h. The stretching membranes were then mounted onto a Strex uniaxial cell stretching system (ST-1500, Green Leaf Scientific) fitted to a Marianas 3i spinning disk confocal microscope (Intelligent Imaging Innovations, Inc). Initial imaging was conducted on cells in their native state using a CMOSS camera (ORCA-Fusion, Hamamatsu), with two channels selected. One channel was used to detect a 544-690nm wavelength range under 561nm excitation with a 300ms exposure, and another to detect a 515-569nm range under 515nm excitation with a 300ms exposure. Cells were then exposed to motor actuation based uniaxial stretching set at a 5% strain for a duration of 2.5 minutes, putting the cells under strain. Following stretching, cells were returned to their unstretched positions and re-imaged, before being re-stretched for another 2.5 minutes. This was cyclically repeated until cells had been stretched for a total of 10 minutes. Between each stretch interval, single z-plane images were captured as previously described using a αPlan-Apochromat 40x/1.4NA oil objective lens (Zeiss).

### Western Blotting

Cells were detached as previously described and transferred into a 1.5 mL Eppendorf tube. The cells were then spun down via centrifugation at 10,000 RPM for 5 minutes and the excess cell culture media was aspirated off. Cells were then resuspended in 100μL RIPA lysis buffer (supplemented with 1x HALT protease and phosphatase inhibitor, Thermo Fisher Scientific, Cat #78440) at 4°C for 15 minutes. The cell lysis solution was then spun down via centrifugation at 12,000 RPM for 15 minutes at 4°C. The supernatant was then transferred into a fresh, pre-cooled 1.5 mL tube and the pellet was discarded. The lysates were then either processed immediately or stored at −20°C until use.

Protein concentration of the lysates was calculated via a Precision Red Advanced Protein Assay. 1 mL of Precision Red reagent (Cytoskeleton, Cat #ADV02-A) was added to a fresh cuvette and used to normalise the readings. 990μL of Precision Red was then mixed with 10μL of cell lysate in a fresh cuvette and incubated at room temperature for 1 minute. The cuvette was then loaded into the spectrophotometer, and the absorbance of the sample was measured at 590nm. This value was then used to calculate the protein concentration within the lysate in μg/mL.

For a singular experiment, unless stated otherwise, all samples were loaded using the same total volume and protein concentration. Generally, 30-50 μg of protein per sample was loaded per well. The calculated volume of cell lysate was mixed with 1x reducing sample buffer and made up to a total volume of 30 μL with RIPA buffer. Proteins were then denatured by heating the samples at 95°C for 5 minutes, before being spun down at 10,000 RPM for 2 minutes. The sample were then loaded into 1.0 mm 12-well pre-cast gels (4-20% SurePAGE^TM^ Bris-Tris Stain-Free Precast Gels, GenScript, Cat #M00652) for protein separation, run using a mini-PROTEAN Tetra Vertical Electrophoresis Cell (BioRad) with 1x Tris-MOPS SDS running buffer (diluted from 20x MOPS-SDS running buffer, Formedium, Cat #MOPS-SDS1000). Electrophoresis was then conducted at 110V for 10 minutes, and then 175V for a further 30-40 minutes.

Proteins were transferred using a Trans-Blot Turbo Transfer System (BioRad) onto a Cytiva Amersham^TM^ Ptotran^TM^ premium (0.45μm pore size) Nitrocellulose membrane (Thermo Fisher Scientific, Cat #10600002). Proteins were transferred for 14 minutes at a constant 1.3Amp and 25V using a 1x Trans-blot Turbo transfer buffer (5x Trans-blot Turbo transfer buffer, BioRad, Cat #10026938) containing 20% (v/v) ethanol.

Transferred proteins were stained using Ponceau stain to check for successful protein transfer, before the transfer membrane was rinsed three times for 5 minutes in TBS-T buffer. The transfer membrane was blocked with 3% BSA blocking buffer (w/v) for 1 hour on a rocker at room temperature. Primary and secondary antibodies were then diluted in 0.3% BSA blocking buffer (in TBS-T) in a 50ml falcon tube. The membrane was then added to the primary antibody tube and rolled at 4°C overnight. After incubation with primary antibodies membranes were washed three times for 5 minutes with TBS-T on a rocker. Blots were then incubated with secondary antibody and rolled at room temperature for 2 hours, before washing the membrane three times in TBS-T for 5 minutes each wash.

For chemiluminescence-cased detection, membranes were incubated in Clarity Western ECL Substrate (BioRad, Cat #170506) following the manufacturer’s instructions. Membranes were then developed and imaged using a Chemidoc^TM^ Touch (BioRad) imaging system for protein visualisation.

### Single cell Migration Assay

6-well cell culture plates (Corning) were coated with 10 μg/mL fibronectin and 50 µg/mL collagen (in PBS) as previously described. 2 x 10^3^ cells were then seeded in the pre-coated 6-wells with DMEM GlutaMax^TM^ media supplemented with 10% FBS and allowed to adhere and spread for 4 hours at 37°C and 5% CO_2_. Cell culture plates were then placed into the live cell incucyte® system (Sartorius, Incucyte® S3 live cell analysis system) and imaged every 30 mins for 48 hours. Images were captured using the built-in HD phase contrast imaging module with a 20x objective. The images were then tracked using FIJI (ImageJ) plug-in TrackMate® (Ershov *et al.,* 2022). 5 cells per panel were tracked until the end of the image stack unless they divided, in which case tracking was stopped at the point of cell division. Total distance travelled (µm) and speed (µm/h) was quantified and plotted. At 0, 12, 24, and 48h lamellipodia area was quantified for each cell. This was identified as the lamellipodia minor axis multiplied by the major axis within the cells leading edge using brightfield images in which the nucleus was clearly visible. Defining the approximate lamellipodia as the area in front of the nucleus and protruding from the front of the cell.

### GFP-Trap Immunoprecipitation

For GFP-Trap® immunoprecipitation (GFP-Trap IP), cells were seeded onto 2 x 10 cm cell culture dishes (Corning) pre-coated with fibronectin (10 µg/mL) and collagen (50 µg/mL) and grown to approximately 80% confluency. Cells were then transfected with either GFP-SKT or GFP-Paxillin, before their lysates were collected 24 h post transfection. Cells were lysed on ice for 30 minutes and centrifuged at max speed for 15 minutes at 4°C. Cell lysates were then transferred to precooled tubes and diluted in immunoprecipitation dilution buffer. 50 µL of was then stored at −20°C as the total protein input fraction, while the remaining samples was incubated with GFP-Trap® agarose beads on rotation at 4°C for 2 hours. The beads were then transferred to new tubes, and the remaining solution was stored at −20°C as the flow through fraction, following magnetic separation. The agarose beads were then washed in immunoprecipitation wash buffer three times and transferred to a new tube during their final wash. Protein complexes were then dissociated from the beads at 95°C with 2x reducing sample buffer for 5 minutes, prior to analysis with western blotting for GFP-Paxillin (as previously described) or with mass spectrometry for GFP-SKT samples.

Samples were separated via polyacrylamide gel electrophoresis. Gels were then stained with instant blue protein dye (abcam) and washed with ddH_2_O. Each sample was then cut into slices and de-stained with repeated 30-minute incubations of 50% (v/v) acetonitrile (ACN)/50% 25 mM NH_4_HCO_3_ (v/v) at room temperature. Gels were then dehydrated with ACN and dried by vacuum, centrifugation. Peptides were then reduced by incubation with 10 mM dithiothreitol (DTT) at 56°C for I hour. Peptides were then alkylated in 55 mM iodoacetamide (IA) in 25 mM NH_4_HCO_3_ for 45 minutes at room temperature in the absence of light. DTT and IA were then removed via washing and dehydration twice as follows: 5 minutes incubations with 25 mM NH_4_HCO_3_, followed by a 5-minute incubation with ACN. The ACN was then removed by centrifugation, and the gel pieces were re-dried by vacuum centrifuge. 1.25 ng/L Porcine trypsin (Promega, Cat #V5280) in 25 mM NH_4_HCO_3_ was then added to the gel pieces and incubated at 4°C for I hour, to allow for trypsin to penetrate the gel before samples were incubated at 37°C for overnight digestion. Trypsinised peptides were then separated from gel pieces by centrifugation and residual peptides were extracted with a 30-minute incubation at room temperature with 99.8% ACN/0.2% (v/v) formic acid, followed by 50%(v/v) ACN/0.1% formic acid (v/v) which were subsequently extracted by centrifugation. The collected eluate was then dried by vacuum centrifuge and peptides were stored at −20°C until resuspension prior to mass spectrometry analysis.

Mass spectrometry was conducted by Professor Patrick Caswell (Caswell lab, University of Manchester, Wellcome Trust Centre for Cell-Matric Research). Dried peptides were resuspended in 5% (v/v) ACN in 0.1% formic acid, before being separated on a Nanoacquity (Waters) Ultra Performance Liquid Chromatography (LC) column coupled to an LTQ-Orbitrap XL (Thermo Fisher) equipped with a nano electrospray source (Proxeon). MS spectra were acquired at a resolution of 30,000 and tandem mass spectrometry (MS/MS) was performed.

### XFe96 Seahorse Analysis

The 96-well seahorse XF pro cell culture plate (Agilent, Cat #103774-100) was coated in fibronectin and collagen as previously stated, and incubated overnight at 4°C. The Seahorse XFe24 sensor cartridge (Aglilent, Lot #W29923) was then hydrated with XF calibrant (pH 7.4) (Agilent, Cat #100840-000) at 37°C in the absence of CO_2_, and 5 x 10^4^ Panc-1 cells were seeded into the seahorse cell culture plate and cultured at 37°C with 5% CO_2_ overnight. 5 wells per conditions were seeded and analysed for all experiments. For the glycolytic stress test: Seahorse XF DMEM medium was then supplemented with 2 mM L-glutamine. For the mitochondrial stress test: Seahorse XF DMEM medium was then supplemented with 2 mM L-glutamine, 1 mM pyruvate, and 25 mM glucose. Culture medium was then removed from cells and cells were washed in seahorse XF assay medium three times, before being incubated in 180 µL seahorse medium at 37°C in the absence of CO_2_ for 1 hour prior to analysis. When analysed with PAA hydrogel substrates, gels were made as previously described, gels were then polymerised in the 96-well seahorse cell culture plates and washed with seahorse XF assay medium.

For the glycolytic stress test (Agilent, Cat #103020-100), glucose, oligomycin, and 2DG were resuspended with the prepared seahorse assay medium as follows:

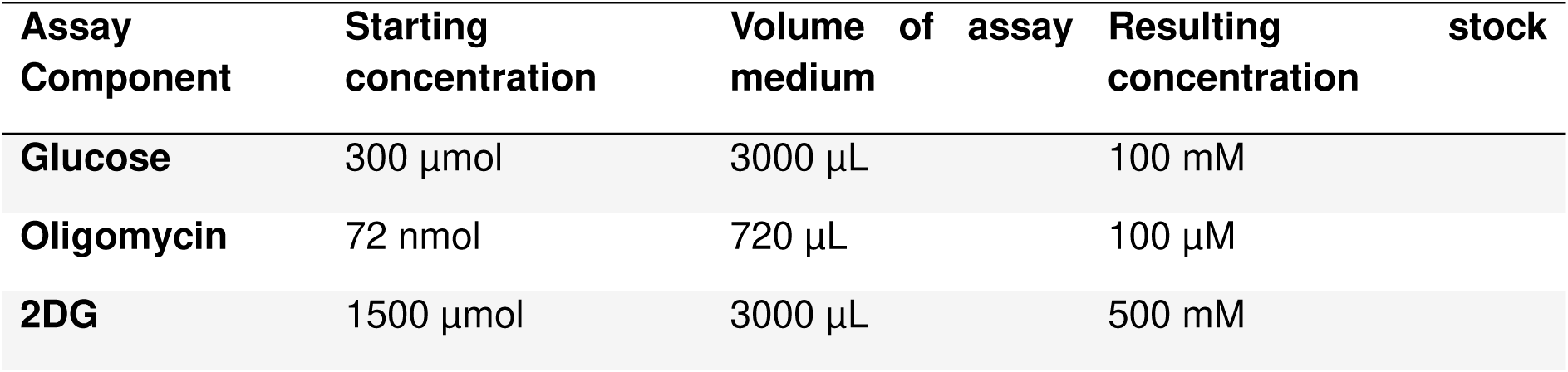

Glycolytic stress test components were then diluted in the prepared seahorse assay medium to the following concentrations: 80 mM glucose, 9 µM Oligomycin, and 500 mM 2DG. 25 µL of each was then loaded into the corresponding injection ports (A – glucose, B – Oligomycin, C – 2DG). The glycolytic stress tests were then ran using the Seahorse XFe24 analyser (Agilent) and extracellular acidification rate was measured for a duration of 100 minutes, in which 4 measurements were taken following each injection. Once injected into the wells, cells were exposed to these components at a final concentration of 10 mM glucose, 1 µM Oligomycin, and 50 mM 2DG.

For the mitochondrial stress test (Agilent, Cat #103015-100), Oligomycin, Carbonyl cyanide-4 (trifluoromethoxy) phenylhydrazone (FCCP), and rotenone and antimycin A (Rot/AA) were resuspended with the prepared seahorse assay medium as follows:

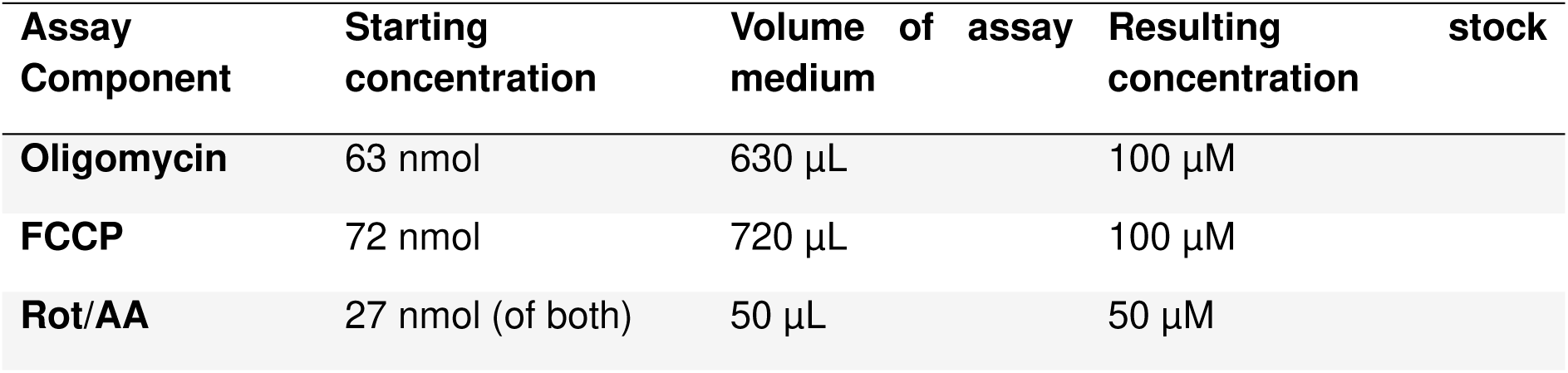

Mitochondrial stress test components were then diluted in the prepared seahorse assay medium to the following concentrations: 10 µM oligomycin, 20 µM FCCP, and 5 µM Rot/AA. 20 µL Oligomycin, 22 µL FCCP, and 25 µL Rot/AA was then loaded into the corresponding injection ports (A – Oligomycin, B – FCCP, and C – Rot/AA). The mitochondrial stress tests were then analysed using the Seahorse XFe24 analyser (Agilent) and oxygen consumption rate was measured for a duration of 80 minutes, in which 3 measurements were taken following each injection. Once injected into the wells, cells were exposed to these components at a final concentration of 1 µM Oligomycin, 2 µM FCCP, and 0.5 µM Rot/AA.

### Promega Glucose-Glo^TM^ Assay

Cells were treated with the appropriate siRNAs as previously described. Following the manufacturer’s instructions (Promega, Cat #J6021), 5 x 10^3^ cells were seeded into 96-well cell culture plate and incubated with Seahorse XF assay media (Agilent, Cat #103680-100) supplemented with 5 mM glucose, L-glutamine, and 10% FBS, and cultured for 48 hours. 2.5 µL of sample medium was then diluted in 397.5 µL PBS and 50 µL was transferred into a new 96-well plate. Glucose detection reagent was then prepared as follows per reaction. On ice,

0.25 µL reductase, reductase substrate, and NAD was added to 50 µL of luciferin detection solution. 2.0 µL glucose dehydrogenase was then added and the mixture was mixed by inversion five times. The luciferin detection solution was equilibrated at room temperature prior to detection reagent preparation. 50 µL glucose detection reagent was then added to the wells simultaneously using a multichannel pipette and placed on a plate shaker for 1 minute. The plate was then incubated for 1 hour at room temperature, before luminescence was analysed using a Glo-Max^TM^ plate reader with an integration time of 0.3 seconds. To calculate glucose concentration, a standard curve was produced using titrations of glucose following serial dilutions of 50 µM glucose (**Supplementary Fig. 2**), confirming a linear relationship between luminescence signal and glucose concentration. For glucose assays on the CytoSoft® imaging and cell culture plates, 5 x 10^4^ cells were seeded. For all glucose detection experiments samples were analysed in triplicate and an extra well was seeded per sample and fixed for cell counts by immuno-staining the nucleus with DAPI as previously described.

### Promega Glucose-Uptake^TM^ Assay

Cells were treated with the appropriate siRNAs as previously described. Following the manufacturer’s instructions (Promega, Cat #J1341), 5 x 10^3^ cells were then seeded into 96-well cell culture plate and incubated with Seahorse XF assay media (Agilent, Cat #103680-100) supplemented with L-glutamine and 10% FBS, overnight. Cell culture medium was then removed, and cells were washed three times in PBS. 50 µL of 1 mM 2-deoxyglucose (2DG) was then added to the wells and mixed for 30 seconds. Cells were then incubated in 2DG for 15 minutes before 25 µL of stop buffer was added to each well simultaneously using a multichannel pipette to lyse the cells, followed by 25 µL neutralisation buffer. 2-deoxyglucose-6-phosphate (2DG6P) detection reagent was then prepared per reaction as follows. On ice, 1 µL NADP^+^, 2.5 µL glucose-6-phosphate dehydrogenase, 0.5 µL reductase, and 0.1 µL reductase substrate was added to 100 µL luciferase reagent. The 2DG6P detection solution was then mixed by inversion and incubated at room temperature for 1 hour prior to use. The luciferin detection solution, stop buffer, and neutralisation buffer was equilibrated at room temperature prior to detection reagent preparation. 100 µL of 2DG6P detection solution was then added to the samples and incubated at room temperature for 2 hours, before luminescence was analysed using a Glo-Max^TM^ plate reader with an integration time of 0.3 seconds. For all glucose uptake assays samples were analysed in triplicate. An extra well was seeded per sample and fixed for cell counts by immuno-staining the nucleus with DAPI as previously described.

### Chick Embryo Model

Fertilised Bovan Brown eggs (Henry Stewart Co., Ltd., Fakenham, UK) were incubated at 37°C and 45% humidity (embryonic day 0; E0) in a specialised poultry incubator (Brinsea OvaEasy 380 Advance EX Series II Automatic Egg Incubator). Eggs were laid horizontally in incubation trays (Brinsea, Weston-super-Mare, UK), and the upward facing side marked to indicate the location for the window to be cut. For the duration of E0–E3, the incubator shelves were set to alternate tilting 45 degrees every 45 min. For in ovo experiments, E3 eggs were windowed by puncturing the wide base of the egg (air cell) with an egg piercer to remove about 5 mL of albumen with a 23 G needle, before sealing the hole with Nev’s Ink tape. Another hole was pierced on the labelled side of the egg, a 3 cm piece of 25 mm 3 M Scotch Magic invisible tape applied, and sharp scissor tips inserted into the pierced hole to carefully cut three sides (2 cm × 1 cm × 2 cm) of a rectangle to create a window in the eggshell. The fenestration area was sealed with approximately 4 cm of 25 mm 3 M Scotch Magic invisible tape, leaving a small tab to enable re-opening of the window, and eggs were placed back into the incubator until E7.

For ex ovo experiments, eggs were gently cracked at E3 and the contents transferred to UV-sterilised black weighing boats (Starlab, Hamburg, Germany). These were placed inside sterile 150 cm^2^ tissue-culture flasks with re-closable lids (Techno Plastic Products AG, Trasadingen, Switzerland) containing sterile water to maintain humidity and incubated until E7 in a Brinsea OvaEasy 360 incubator.

Prior to implantation on E7, GFP-SKT and GFP control cells were collected by trypsinisation, counted, washed in sterile PBS, and pelleted via centrifugation. Experiments were optimised by implanting between 0.5 x 10^6^ and 2 x 10^6^ cells per egg in 10-15 µl. Prior to adding the cells, the chorioallantoic membrane (CAM) was traumatised using a 1 cm wide strip of sterile gauze swab. The cells were directly pipetted on this region of the CAM. The eggs were then resealed and incubated until E14. Chick survival and tumour progression were monitored during experiments, with bioluminescence imaging (BLI) performed prior to fluorescence imaging and tumour dissection at E14.

Following BLI at E14, tumours were imaged under a Leica M165FC fluorescence stereomicroscope with 16.5:1 zoom optics, fitted with a Leica DFC425 C camera (Leica Biosystems, Wetzlar, Germany). Tumours were imaged in ovo on the CAM and then dissected. Briefly, extra-fine straight-tip tweezers were utilised to manipulate the CAM to allow a sizeable circumference to be cut around the tumour nodules with spring bow micro scissors. Excised tumour nodules were placed in sterile PBS for ex ovo imaging. After removing any excess CAM, dissected tumours were weighed on a fine balance and processed for immunohistochemical or transcriptional analyses. Following removal of the tumour, embryos were terminated in accordance with the UK Animals Scientific Procedures Act 1986 (amended 2012), under which the chick embryo is classified as non-protected until two thirds of gestation is reached at E14. No home office approval is required, and procedures were reviewed by the Liverpool Animal Welfare and Ethical Review Body.

### Statistical Analysis

All statistical analysis was conducted using GraphPad (Prism 10.2.1). standard deviations were monitored using descriptive statistics function and Gaussian distribution was tested using the normality and lognormality test’s function, in which a Shapiro-Wilk normality test was used. For all experiments comparing the means of two groups, an independent t-test was used for statistical analysis. If analysis was used to compare three or more groups a one-way ANOVA was conducted. Additionally, if the data contained three or more groups and/or compared the means between two factors, a two-way ANOVA was used. If the data did not display a gaussian distributions, then statistical tests were corrected with non-parametric test. For t-tests, this included a Mann-Whitney test to compare ranks, and for one-way ANOVA this included a Kruskal-Wallis test. Significances were indicated in the figures with the level of significance and test details included in the figure legends.

## Acknowledgments

We thank the University of Liverpool Centre for Cellular Imaging (CCI) (MRC [Medical Research Council] grants BB/T017813/1 and MR/X013502/1) for their support and access to equipment. The Authors acknowledge use of the Egg Facility (RRID:SCR_026195) provided by Liverpool Shared Research Facilities, Faculty of Health and Life Sciences, University of Liverpool. M.C was funded by Biotechnology and Biological Sciences Research Council doctoral training program (BB/M011186/1/1797330). P.C. was funded by the Wellcome Trust Centre for Cell-Matrix Research by grant 203128/Z/16/Z. D.N. was funded by Wellcome Trust PhD studentships (105350/Z/14/A). E. G was funded by MRC project grant MR/Y001125/1 (to T.Z.). P.A. is grateful for start-up funding from the Royal Society (RG\R1\241403). B.H. was funded by a Discovery Medicine North (DiMeN) Medical Research Council doctoral training program scholarship. We would like to thank Dr Mark Morgan and his lab members for advice with the traction force microscopy experiments and Dr. Amy Chadwick for training and assistance on the SEAHORSE system. We would like to thank ADDGENE and the generous laboratories that have made the following plasmids available that we used in this study mAPPLE-Paxillin-22 (Addgene #54935 from Michael Davidson lab); pRK5-YFP-RICTOR (Addgene #73387 from Jie Chen/ Taekjip Ha labs).

---

**Supplementary Figure 1.**
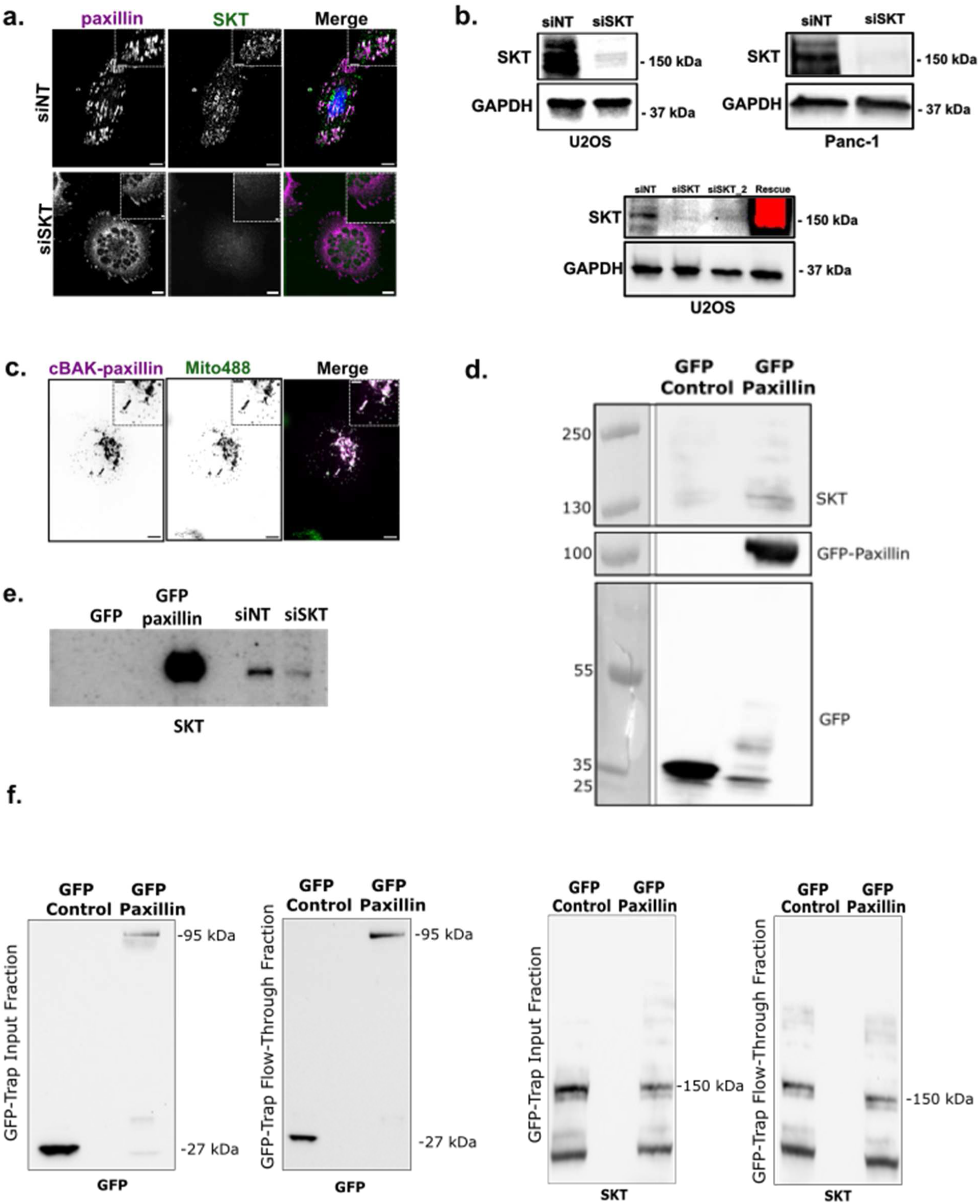
**a.** Representative images of U2OS cells treated with non-targeting control siRNA (siNT) and SKT-targeting siRNA (siSKT) and immuno-stained for endogenous paxillin (magenta) to visualise Integrin-based adhesion complexes (IACs) and endogenous SKT (green). **b.** Representative western blots for U2OS and Panc-1 cells treated with siNT and siSKT, and U2OS cells treated with siNT, two independent SKT targeting siRNAs (siSKT, siSKT_2), and rescued with overexpression of GFP-SKT following treatment with siSKT_2). Blots probed for SKT to assess knockdown efficiency of SKT expression, and GAPDH as a loading control. **c.** Representative images for the mitochondrial targeting assay showing U2OS cells transiently transfected with cBAK-paxillin (magenta), and counter stained with MitoTracker488^TM^ (green) to visualise mitochondria and confirm successful localisation of cBAK-paxillin at the mitochondria. **d.** GFP-Trap immunoprecipitation (IP) of GFP-paxillin. Representative western blot probed for SKT, paxillin, and GFP following IP of U2OS cells expressing GFP-paxillin. The same IP sample was analysed on two separate blots to probe for SKT, and GFP and paxillin. **e.** Representative western blot of GFP-Trap immunoprecipitation (IP) of GFP-paxillin showing lysates from cells transfected with GFP control and GFP-Paxillin, ran alongside lysate from U2OS cells treated with siNT and siSKT. Blot probed for SKT to confirm identification of SKT band. **f.** Representative western blot of the full GFP-Trap IP blots probed for GFP and SKT, with both the input lysates and flow through lysate blots probed for paxillin and SKT to confirm successful capture of paxillin and SKT via the GFP-Trap.

**Supplementary Figure 2.**
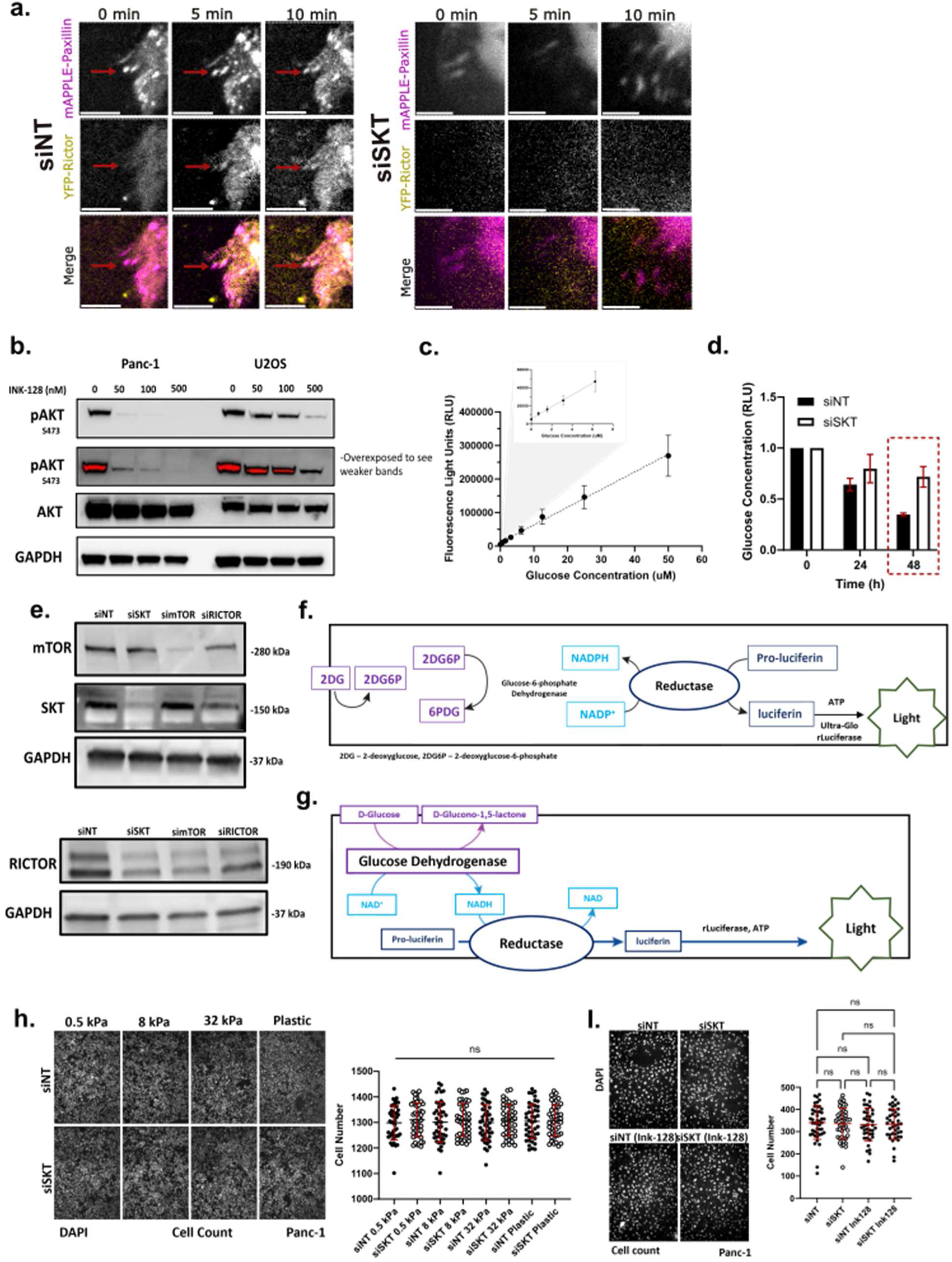
**a.** Representative zoomed inset images of non-targeted siRNA (siNT) and SKT-targeted siRNA (siSKT) treated YFP-Rictor and mAPPLE-paxillin transiently transfected U2OS cells under uniaxial stretching with at 0-, 5-, and 10-min post stretch. **b.** Representative western blots of Panc-1 and U2OS cells treated with 0, 50, 100, and 500 nM dual mTORC1/2 inhibitor INK-128. Blots probed for pAKT and total AKT, and GAPDH as a loading control. **c.** Standard curve of absorbance measurements in relative light units (RLU) using titrations of glucose following serial dilutions of 50 µM glucose to confirm linear relationship between glucose concentration and luminescence signal. **d.** Quantification of glucose consumption in Panc-1 cells at 0, 24, and 48 h. Concentrations normalised to 0 h. **e.** Schematic diagram of the metabolism and reaction pathway utilised in the Glucose-Glo^TM^ assay (Promega) which detects glucose to produce luminescence via ATP-dependent production of luciferin. **f.** Schematic diagram of the metabolism and reaction pathway utilised in the Glucose-Uptake^TM^ assay (Promega) to detect 2-deoxyglucose (2DG) uptake and produce luminescence via ATP-dependent production of luciferin. g. Representative images showing DAPI stained nuclei of siNT and siSKT treated Panc-1 cells on 0.5-, 8-, and 32-kPa substrates with quantification of cell counts post glucose consumption assay. H. Representative images showing DAPI stained nuclei of siNT and siSKT (with and without INK-128 treatment) treated Panc-1 cells, with quantification of cell counts post glucose consumption assay. All results are displayed as mean ± SD per cell. Significance was tested with a two-way ANOVA for mean glucose consumption comparison, ns, P ≥ 0.05.

**Table S1.**
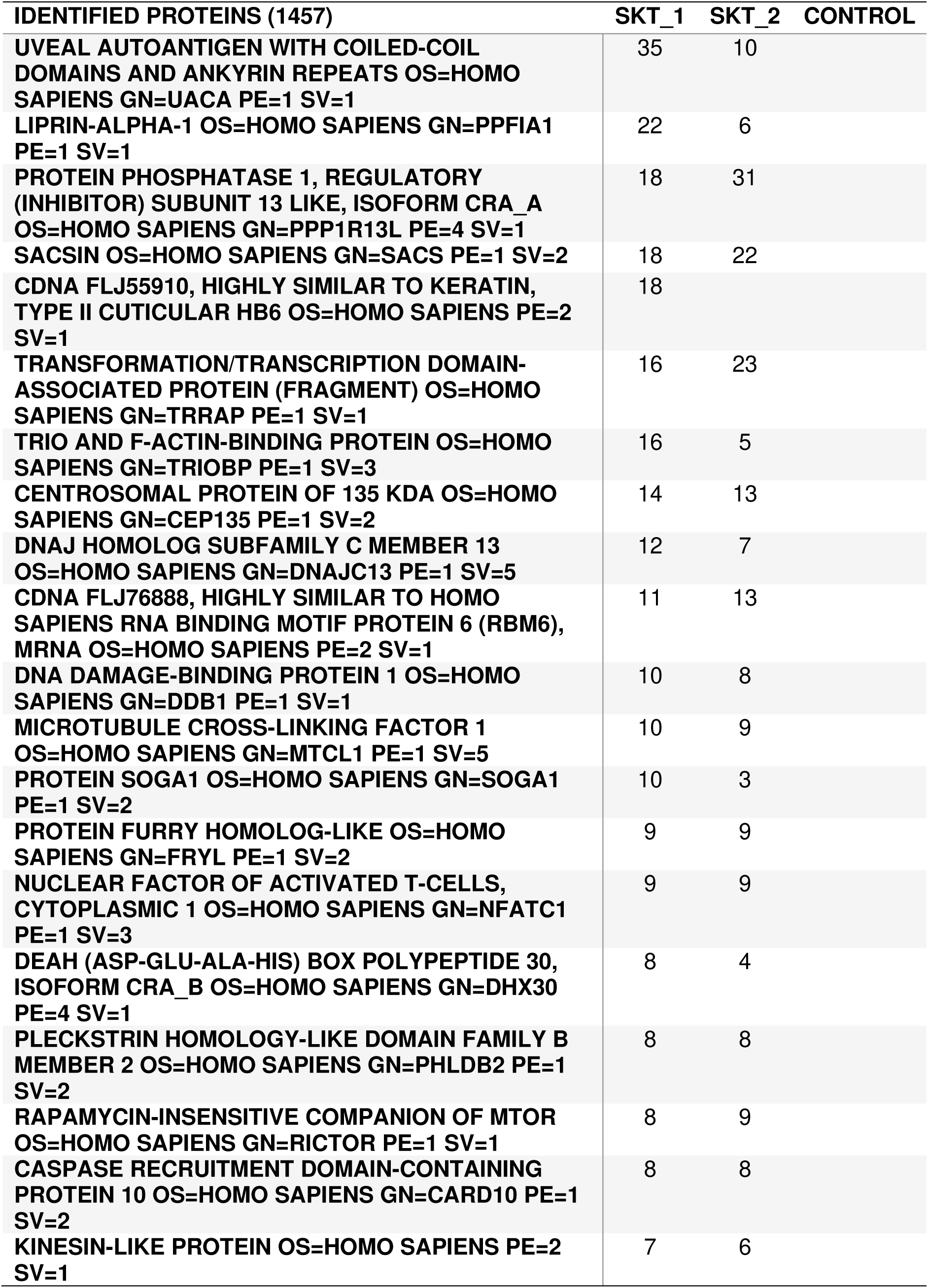
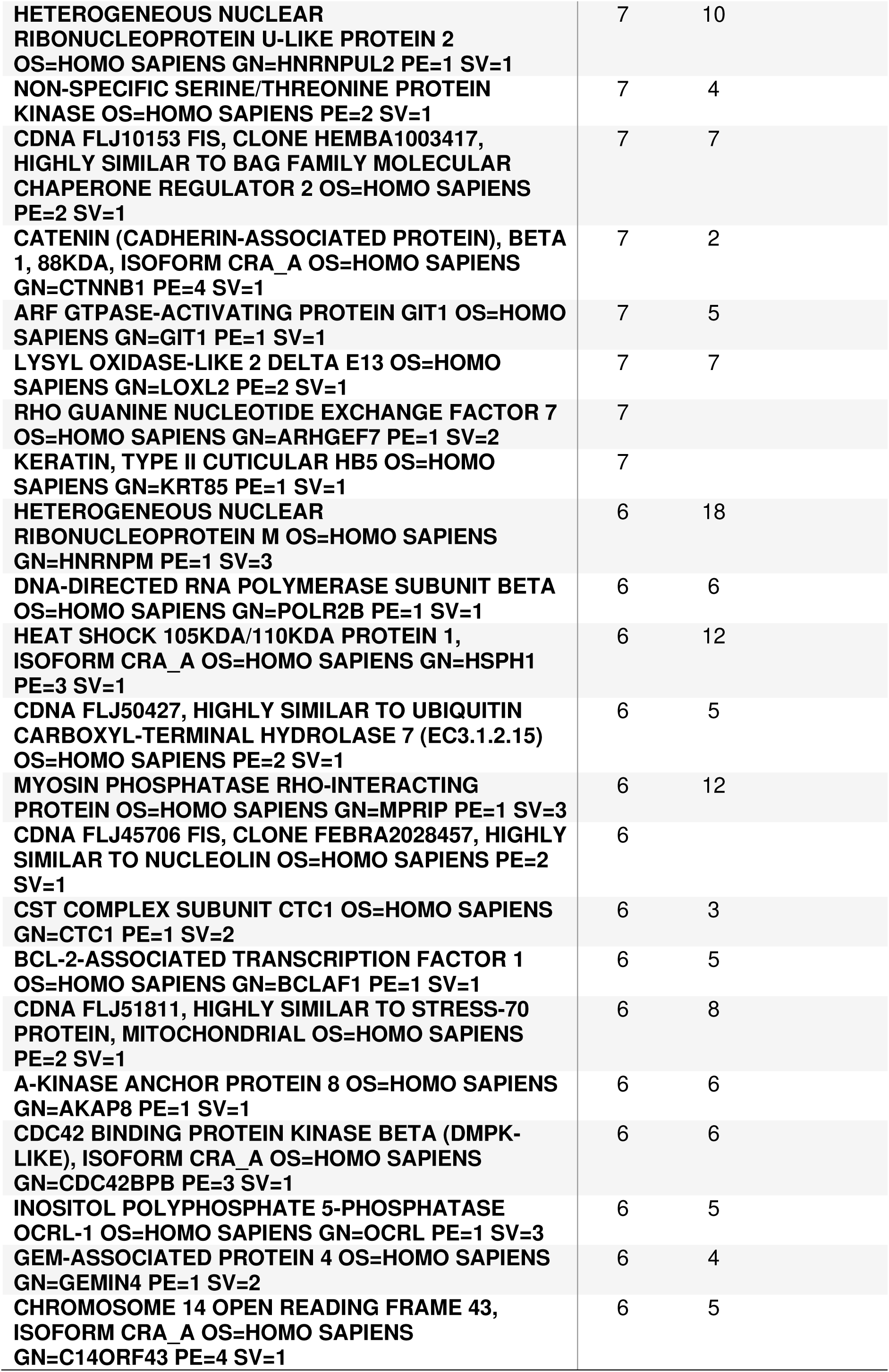
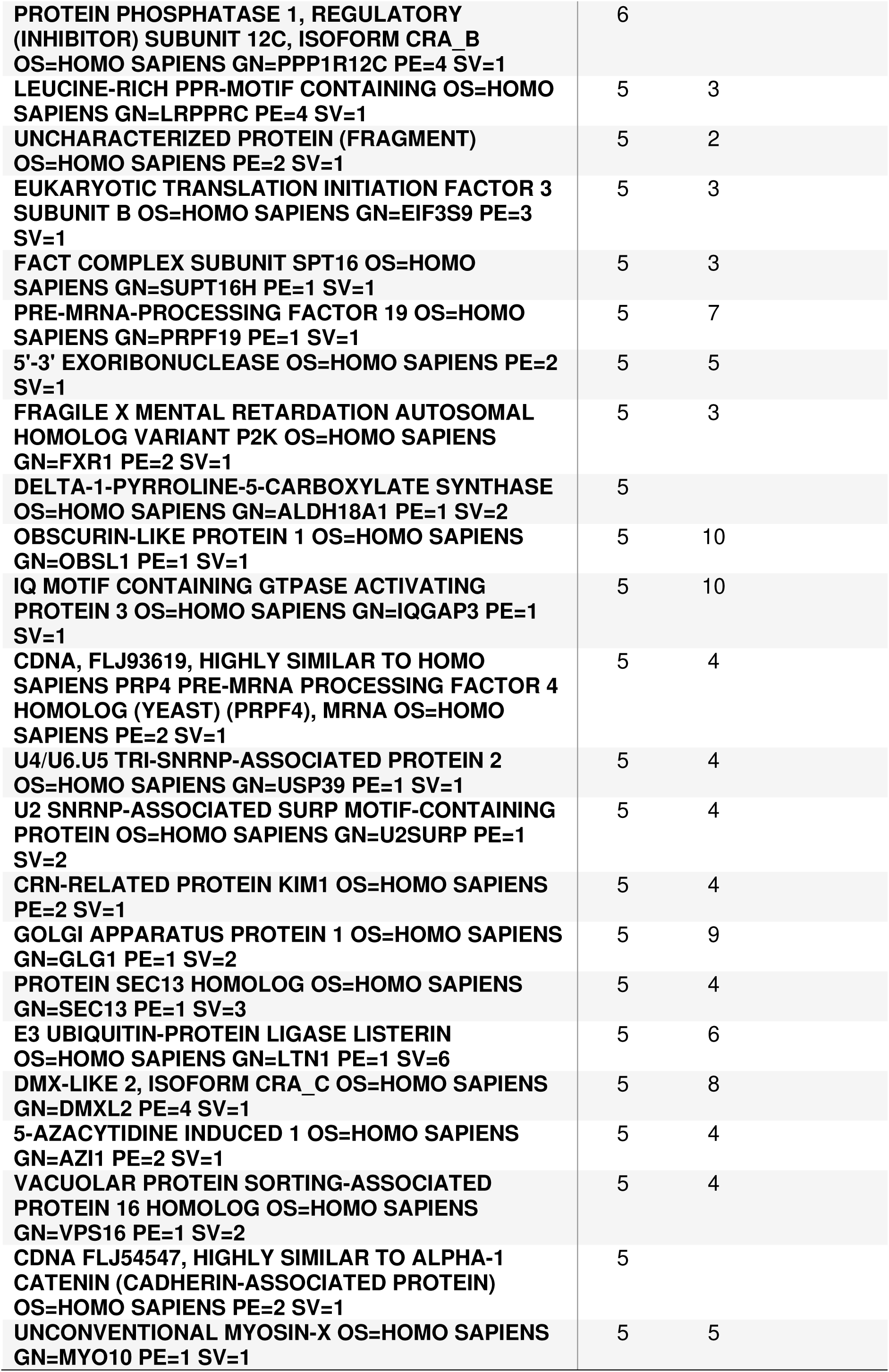
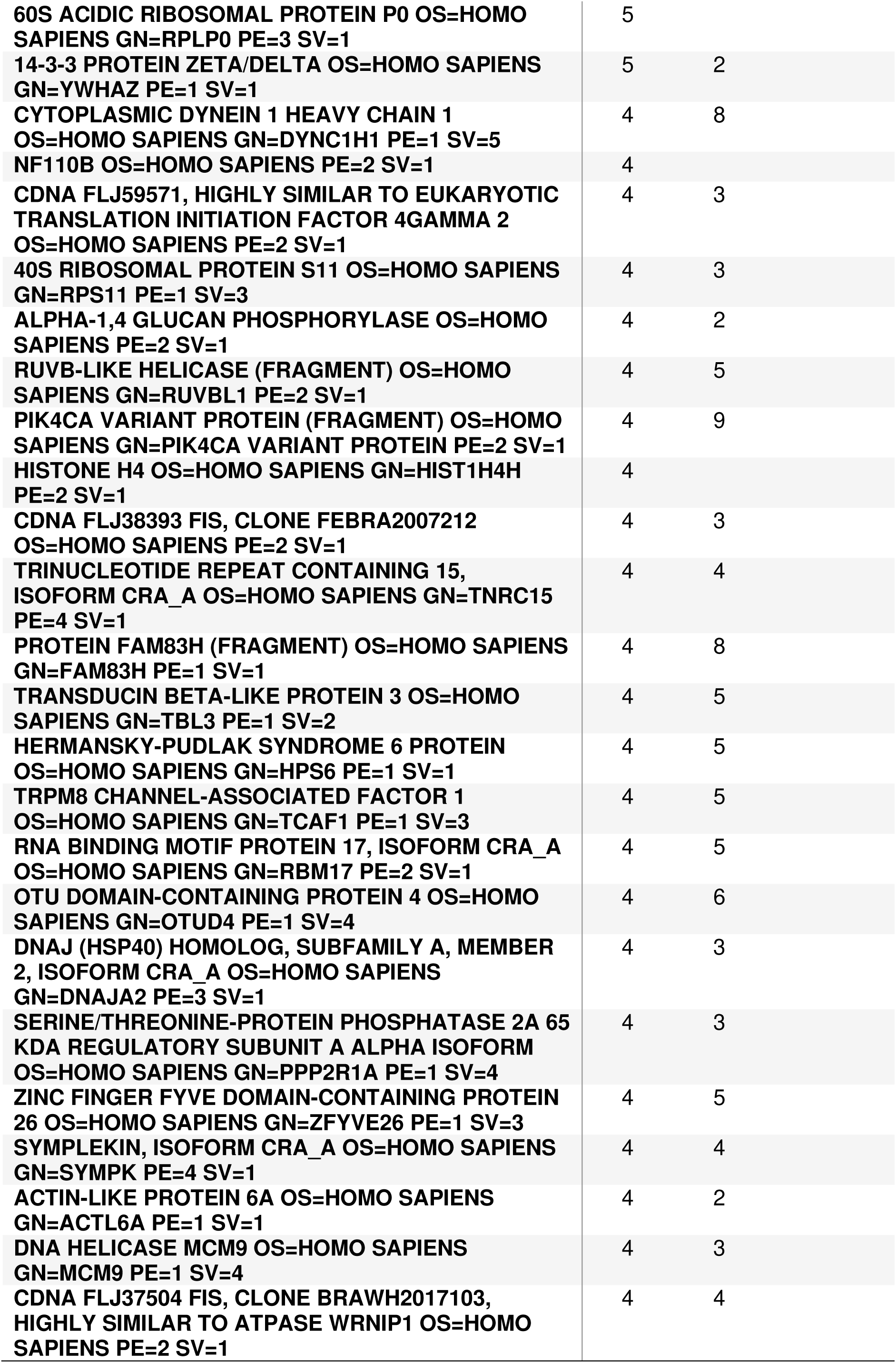
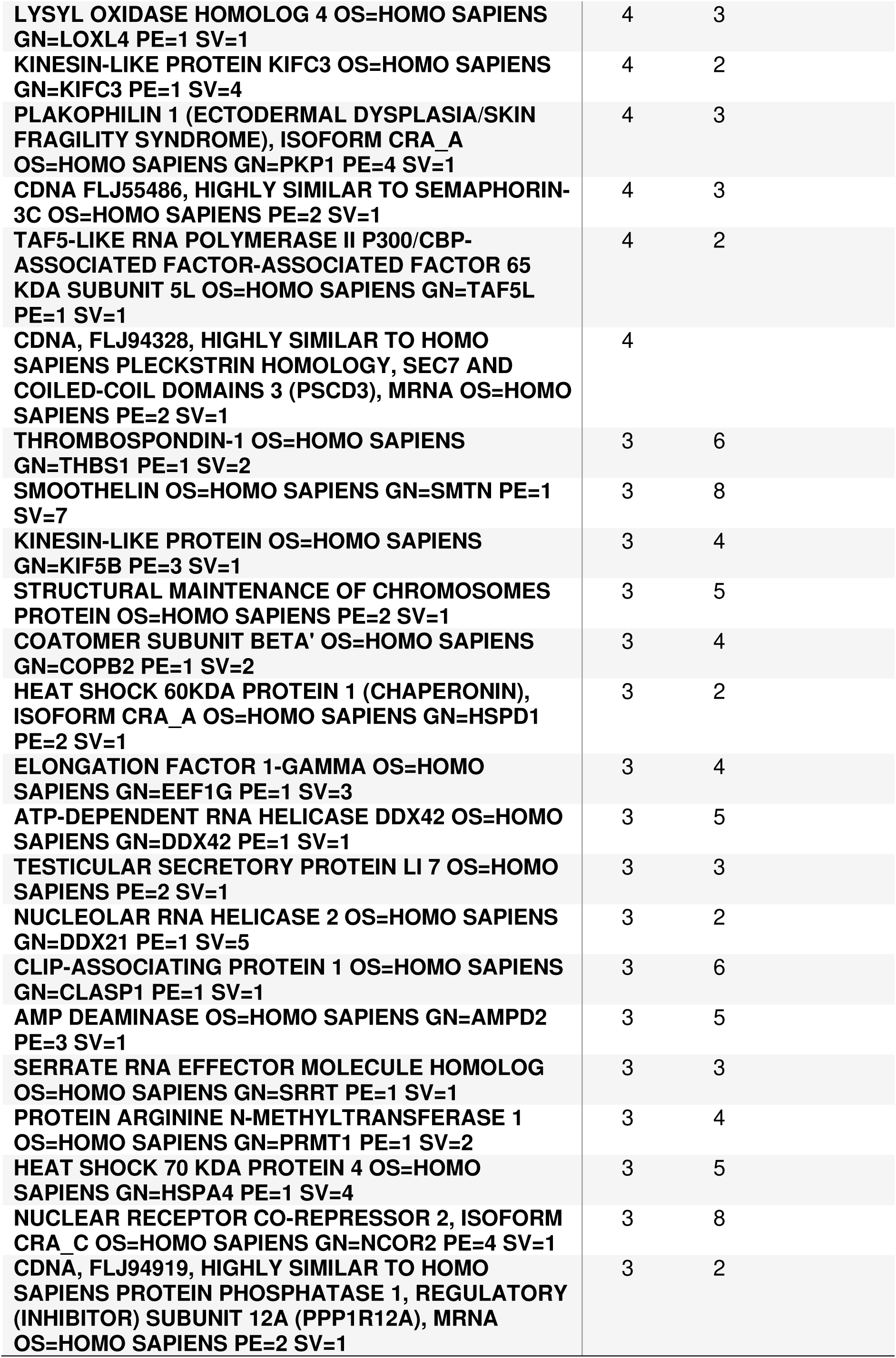
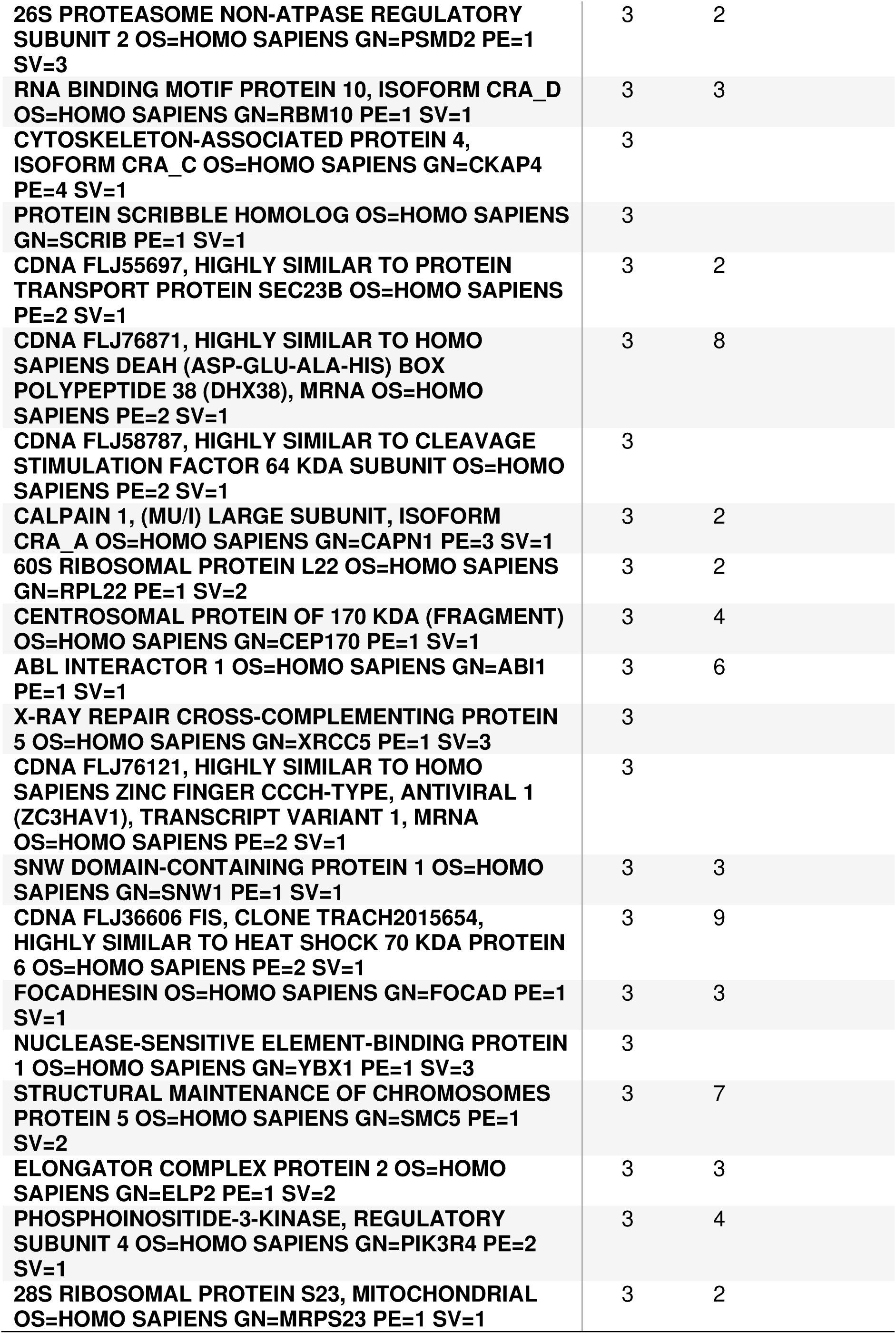
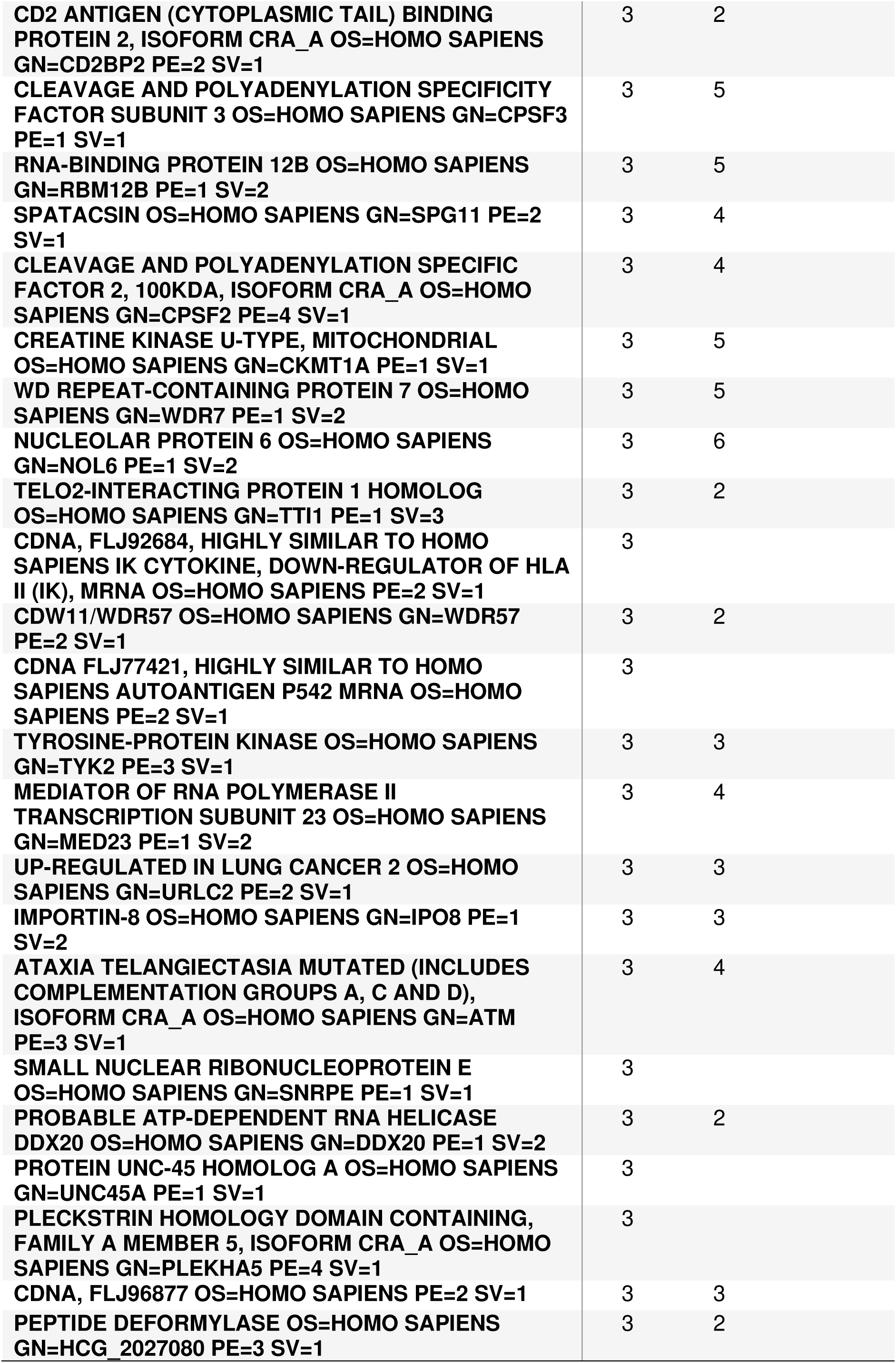
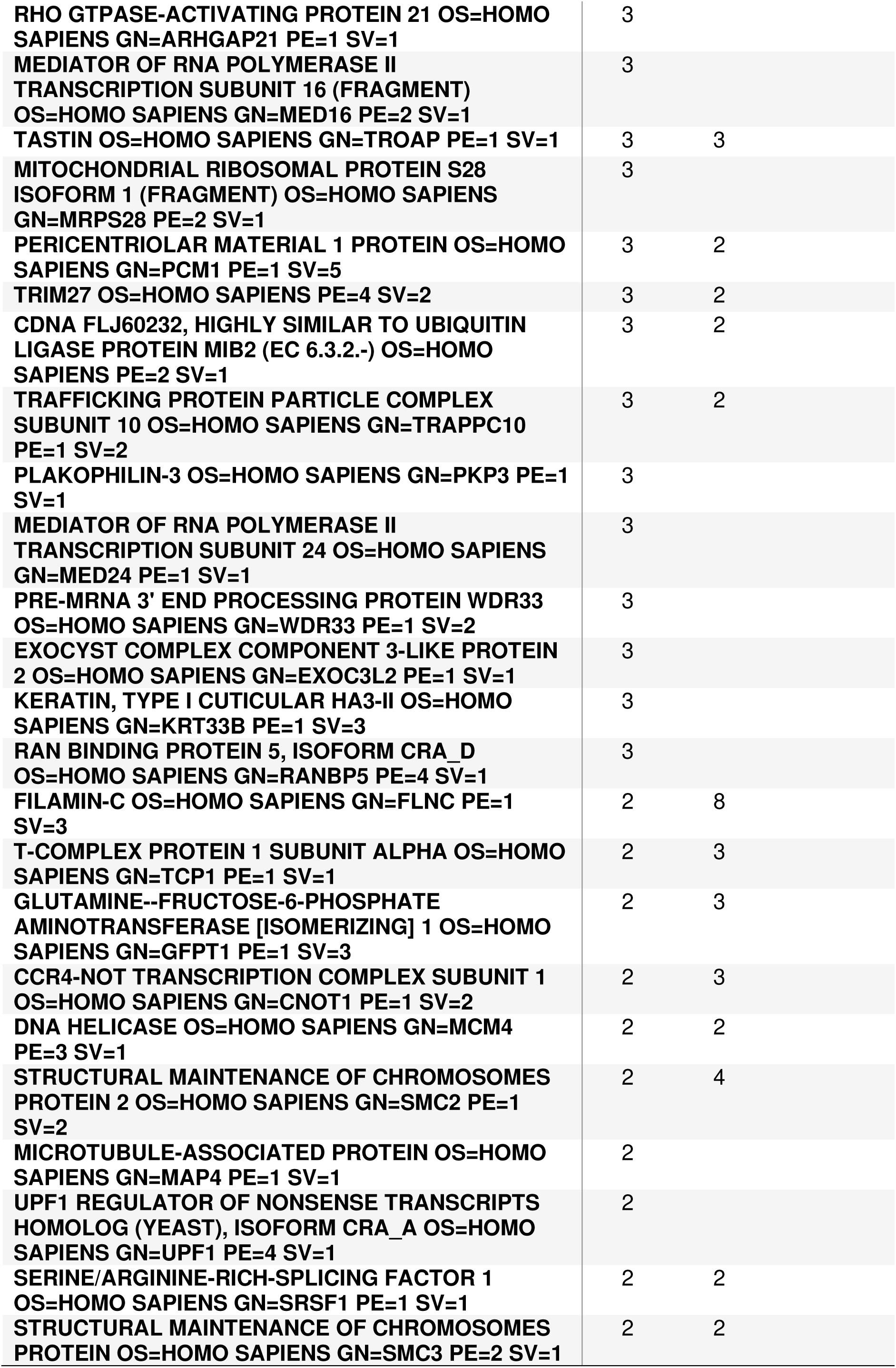
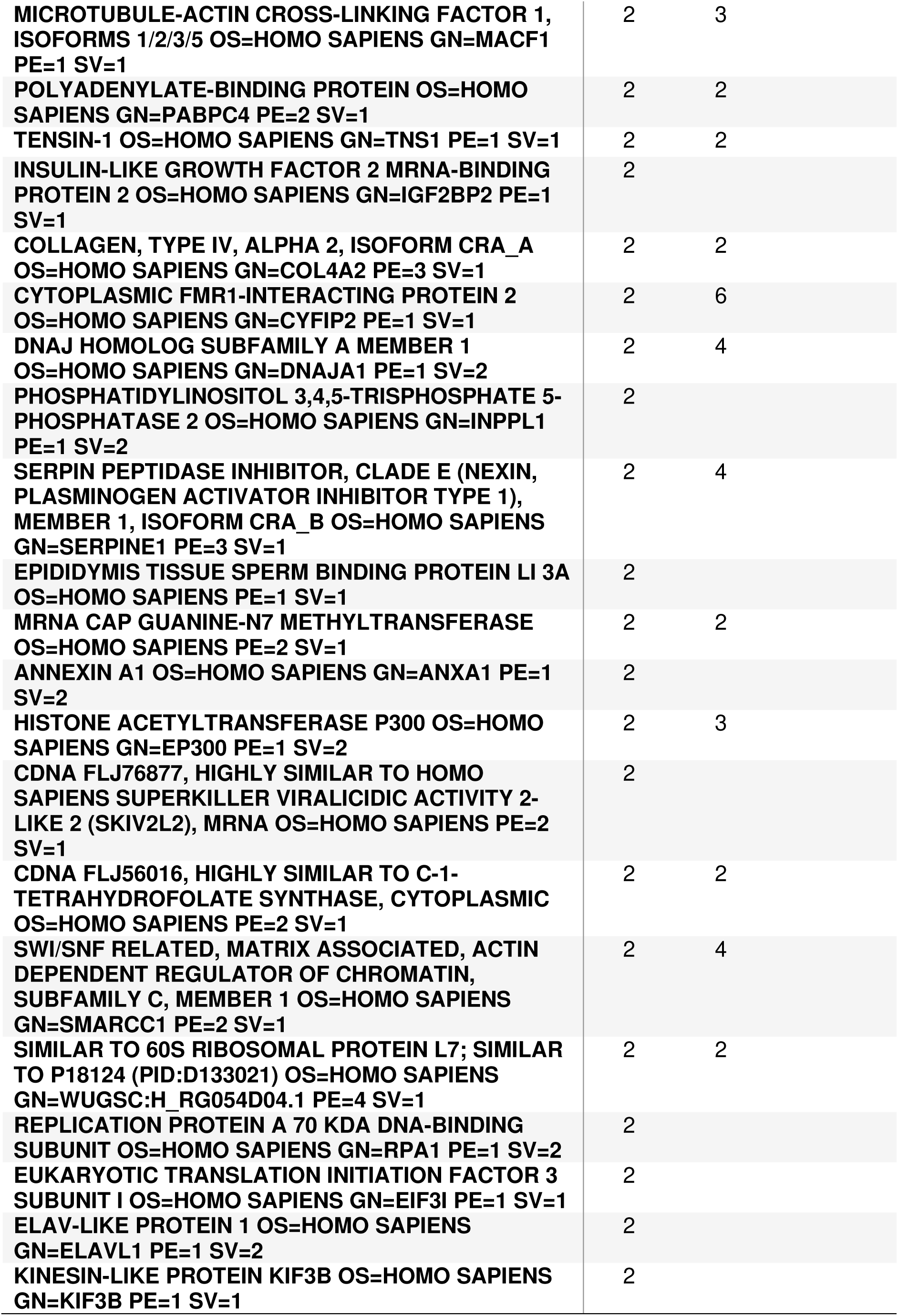
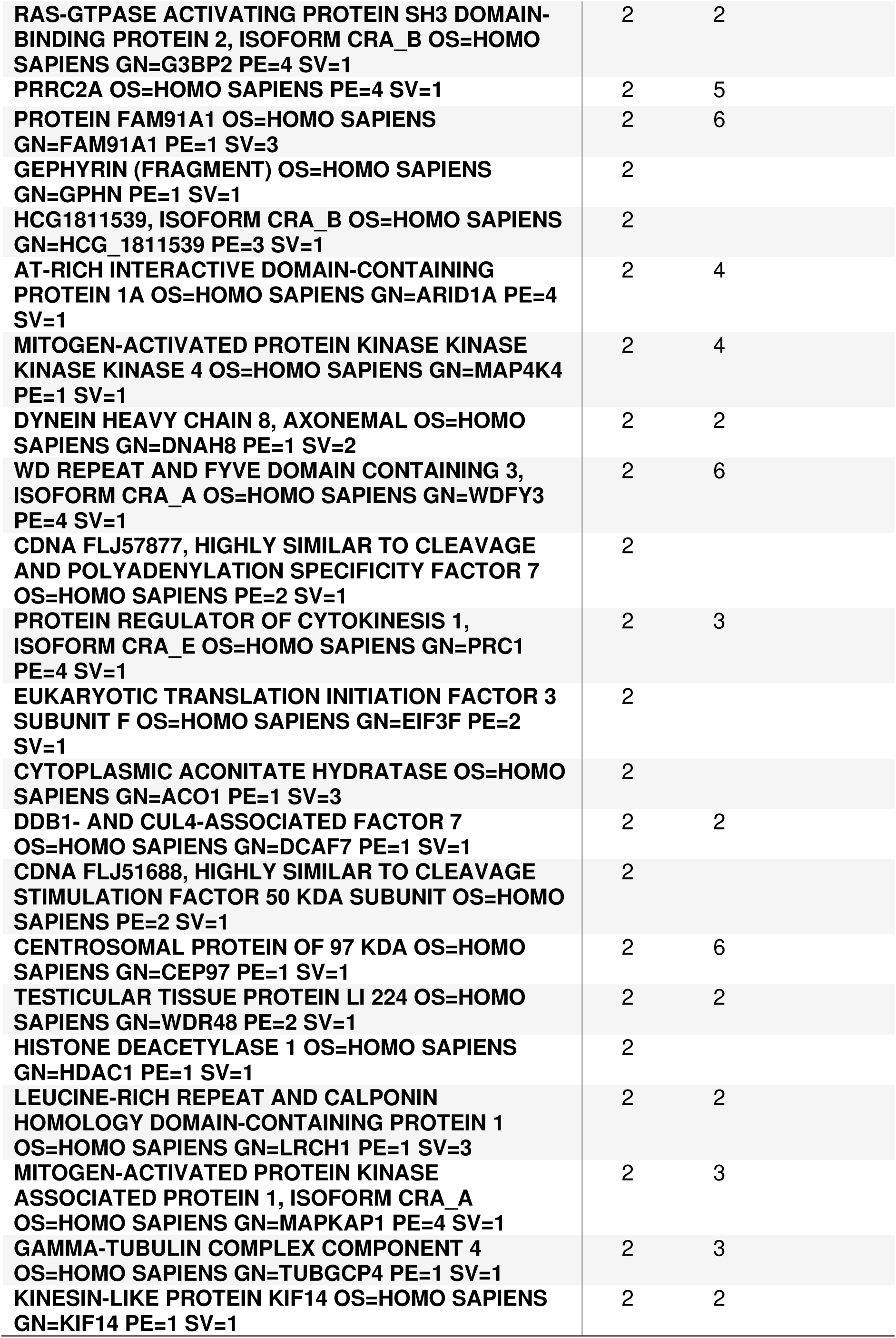
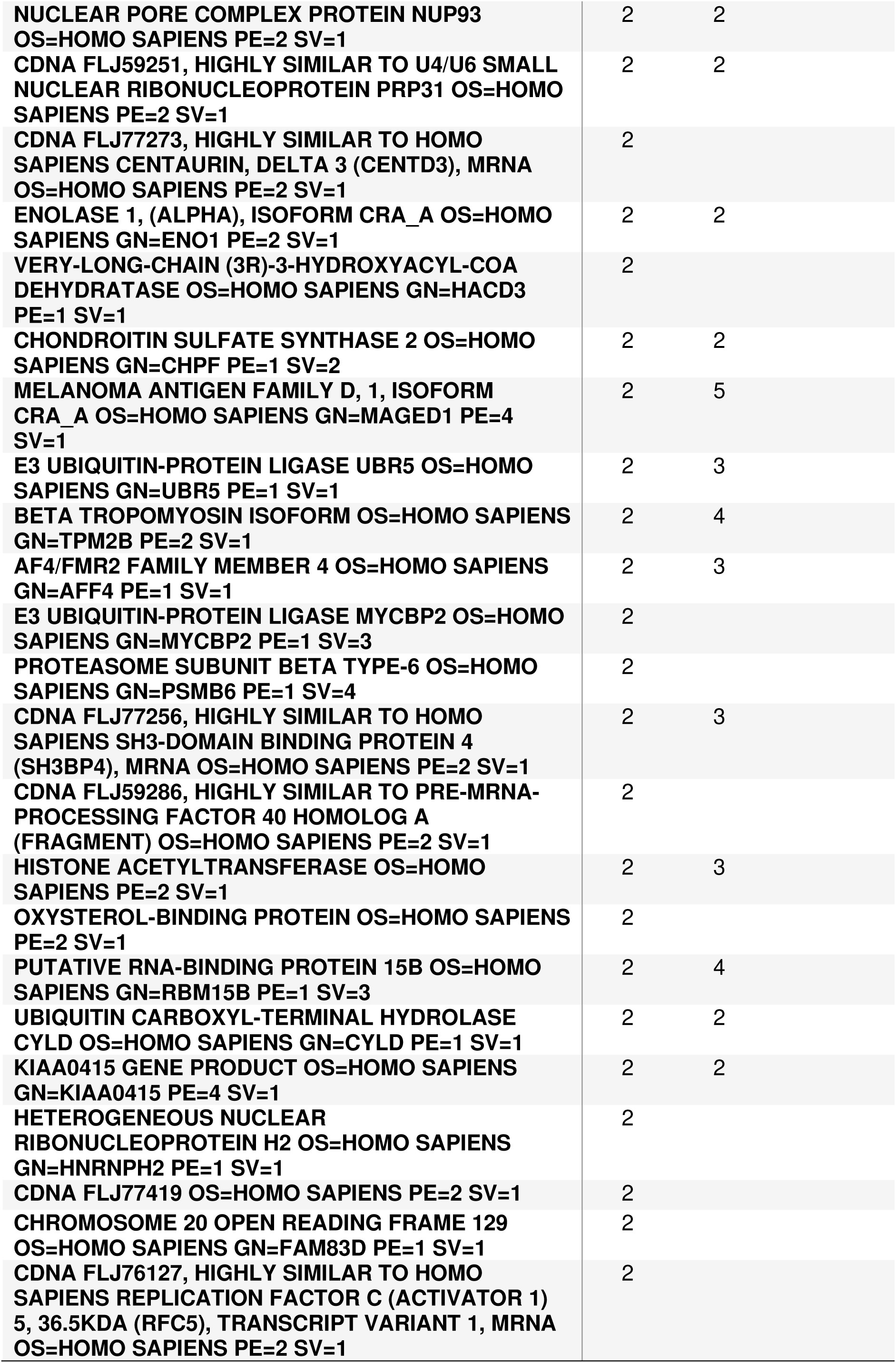
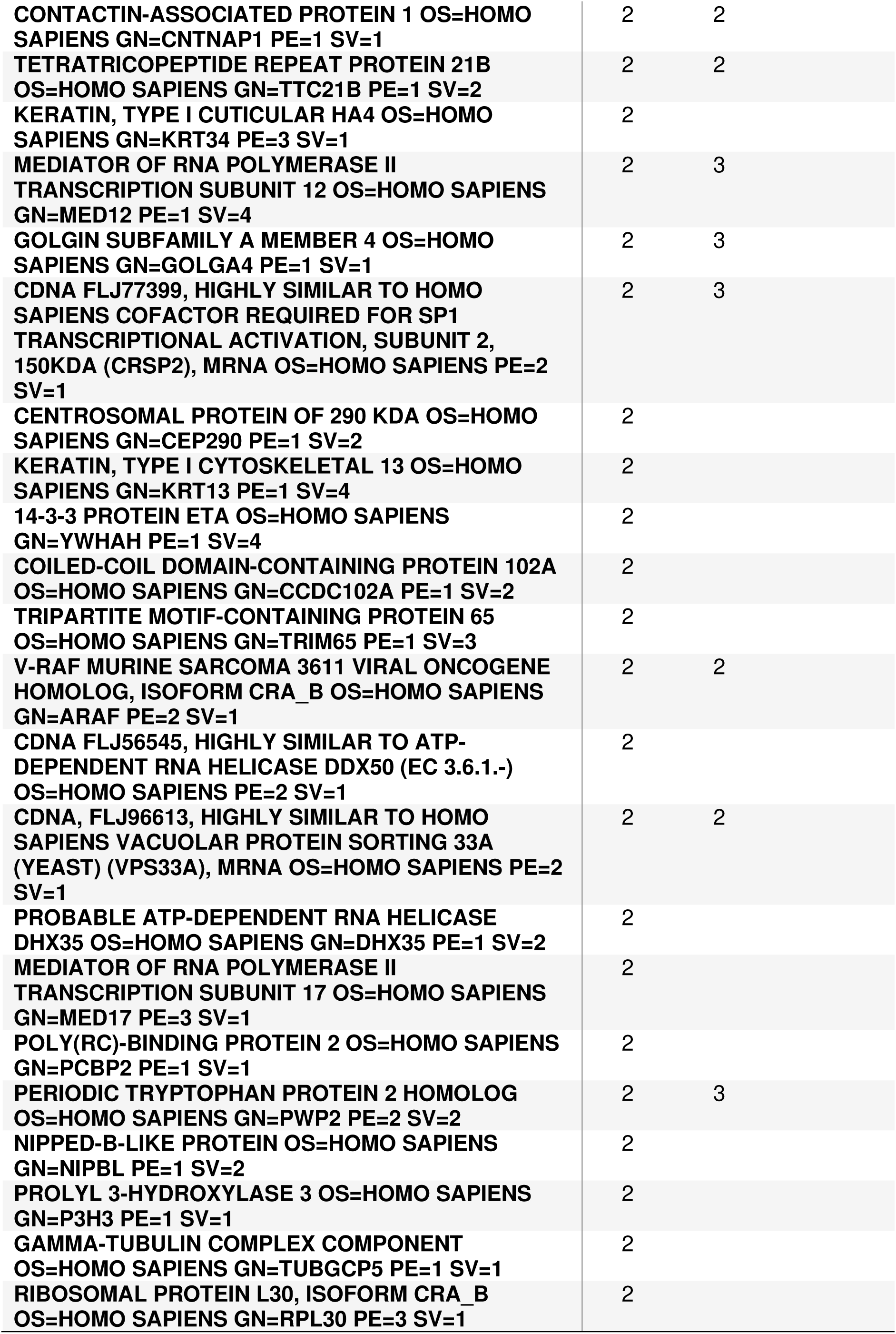
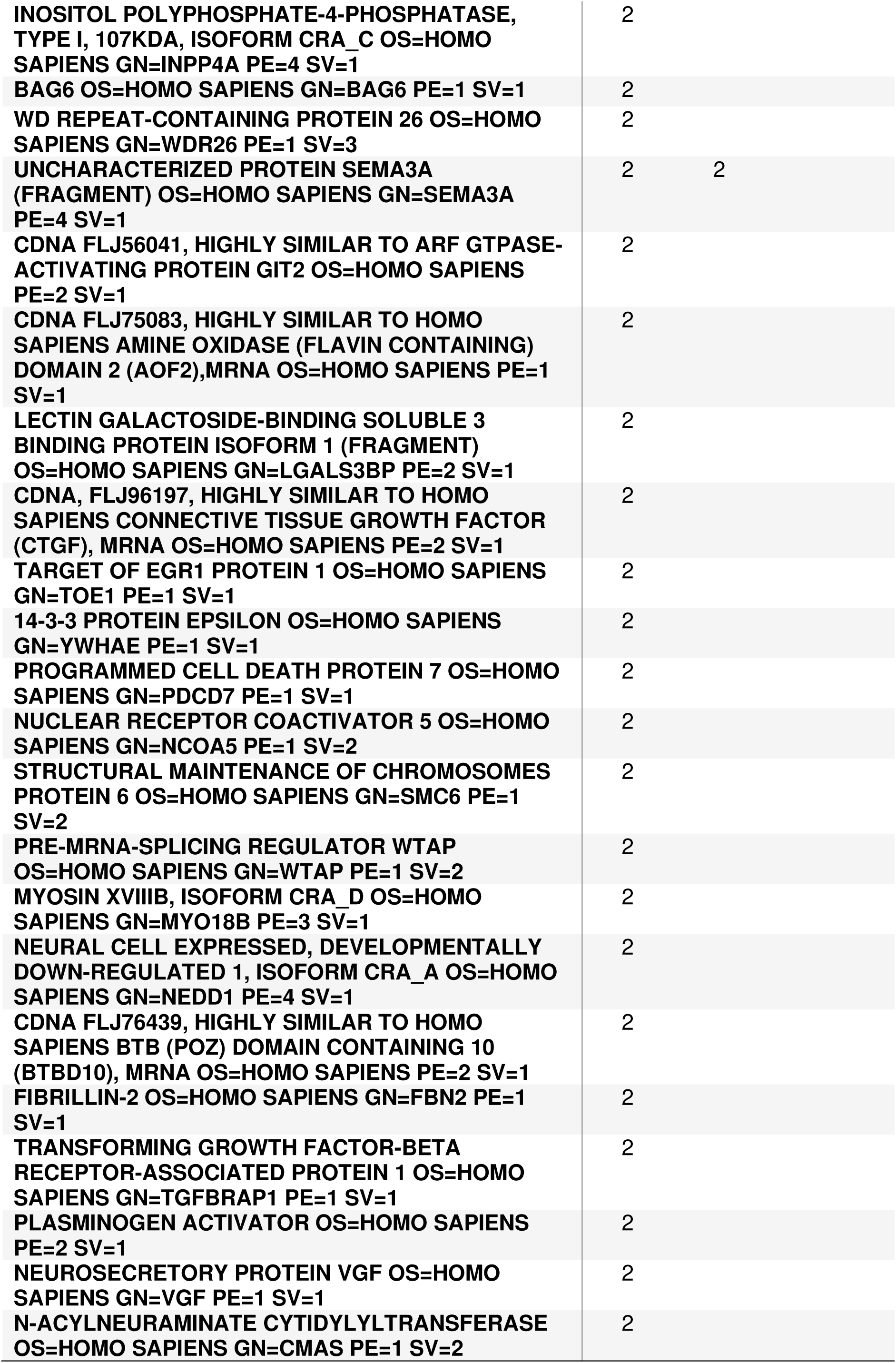
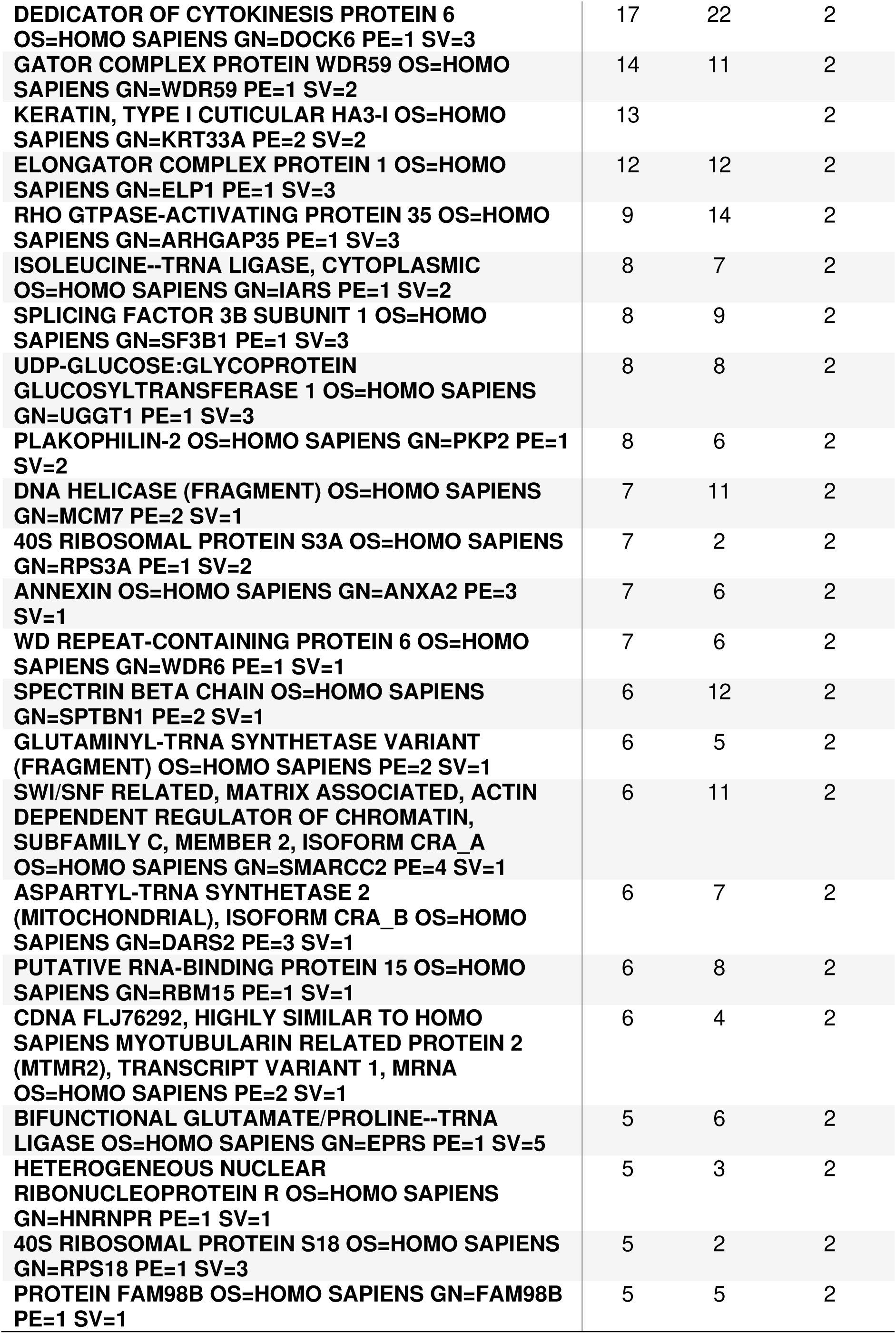
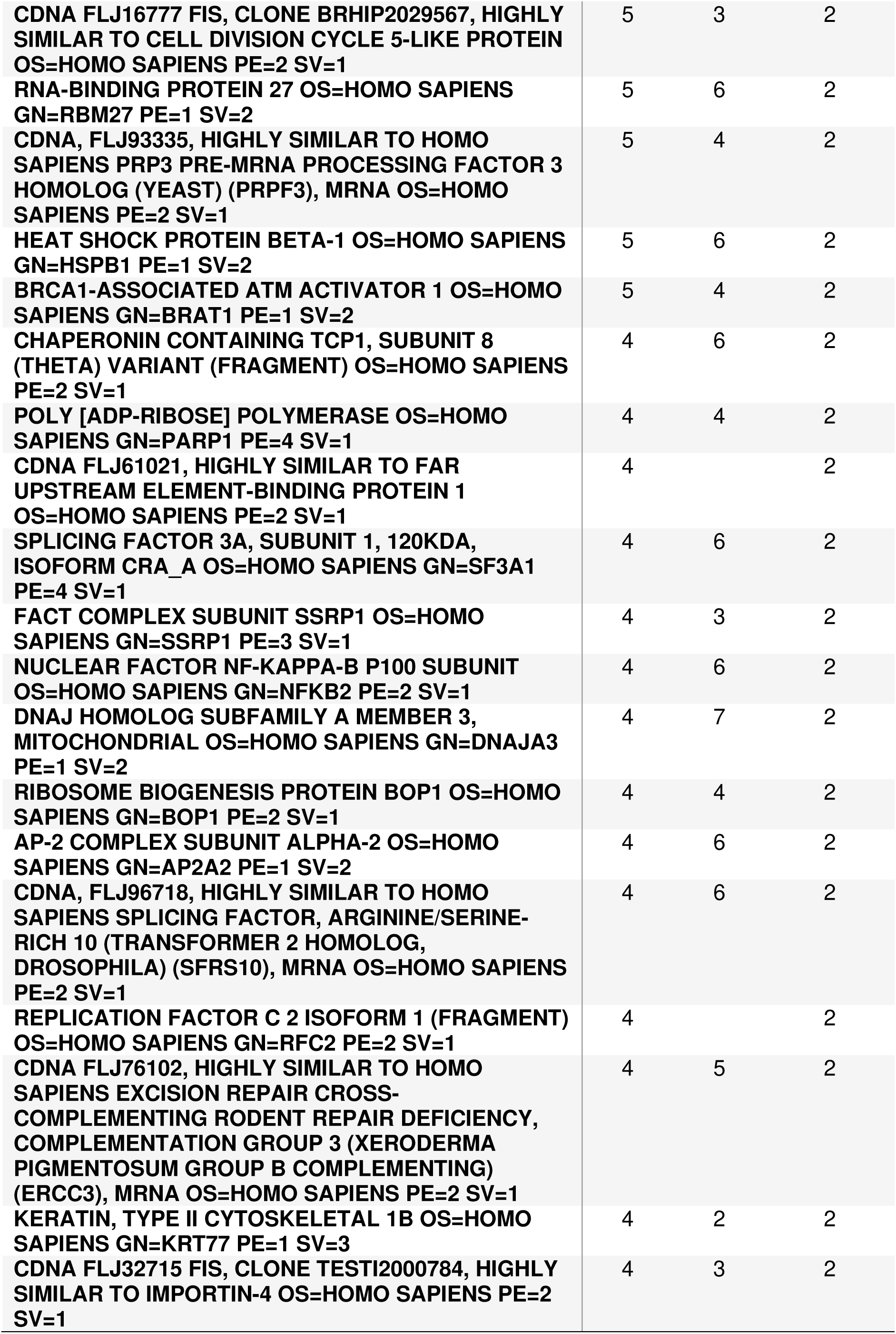
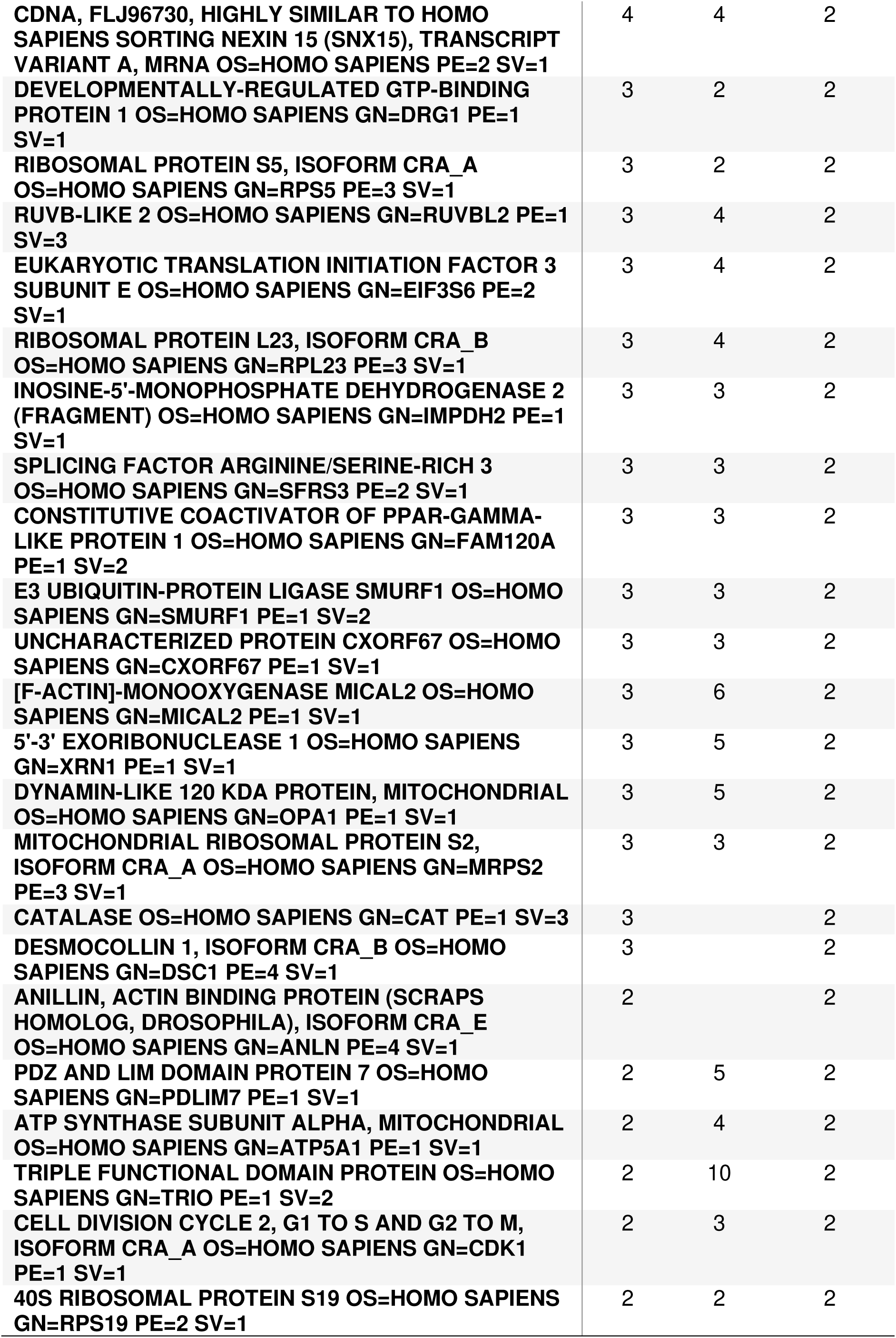
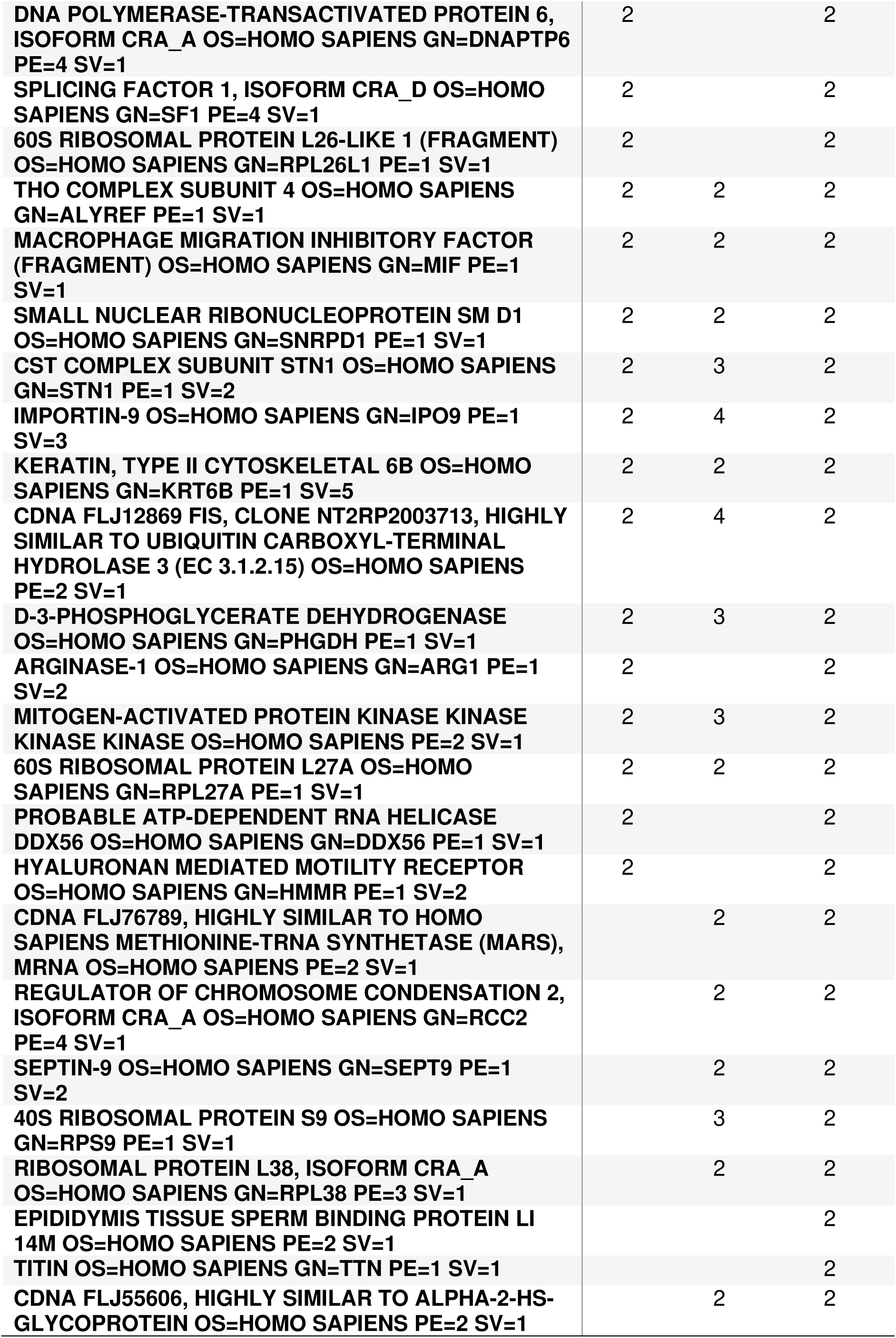
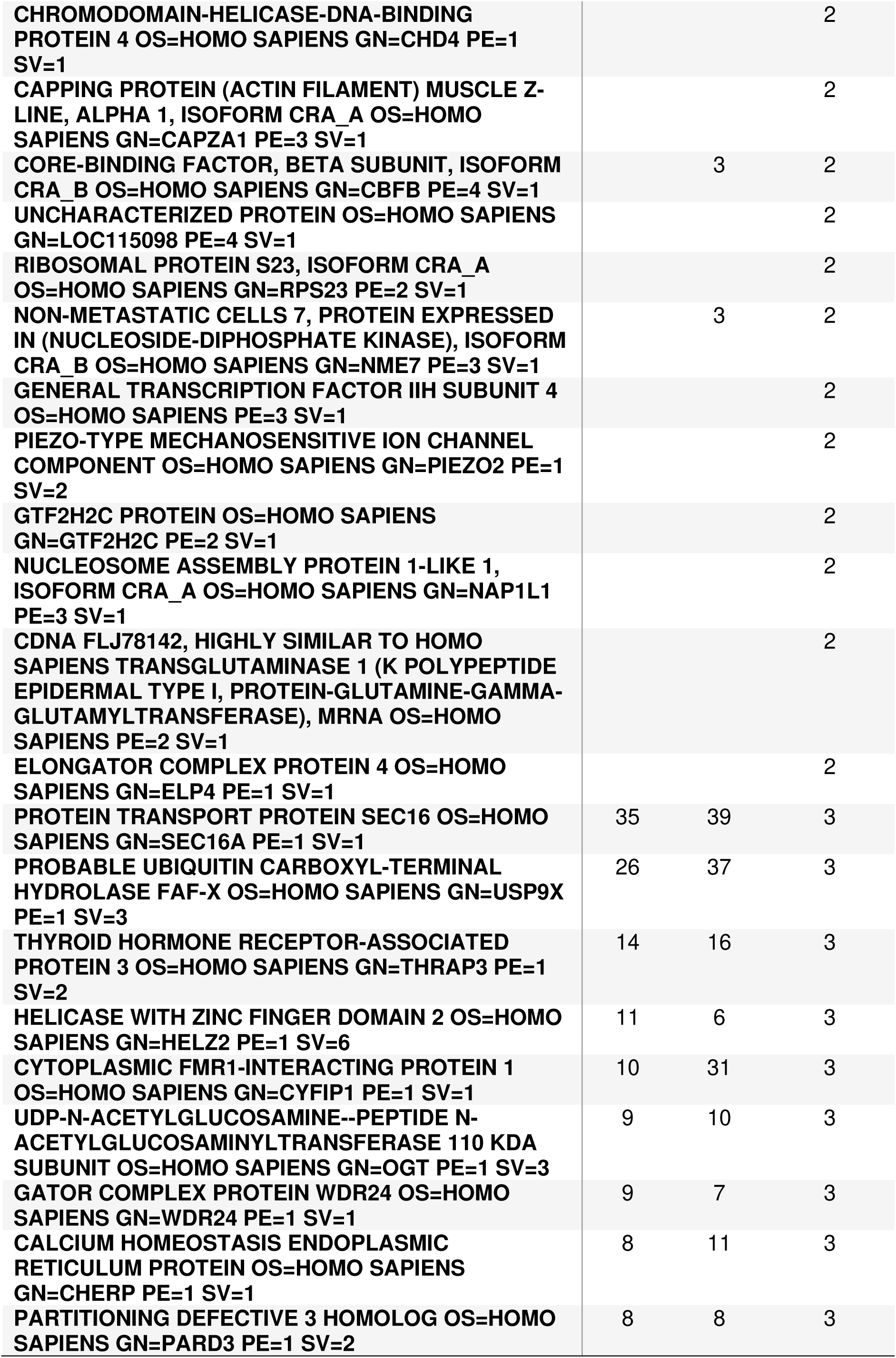
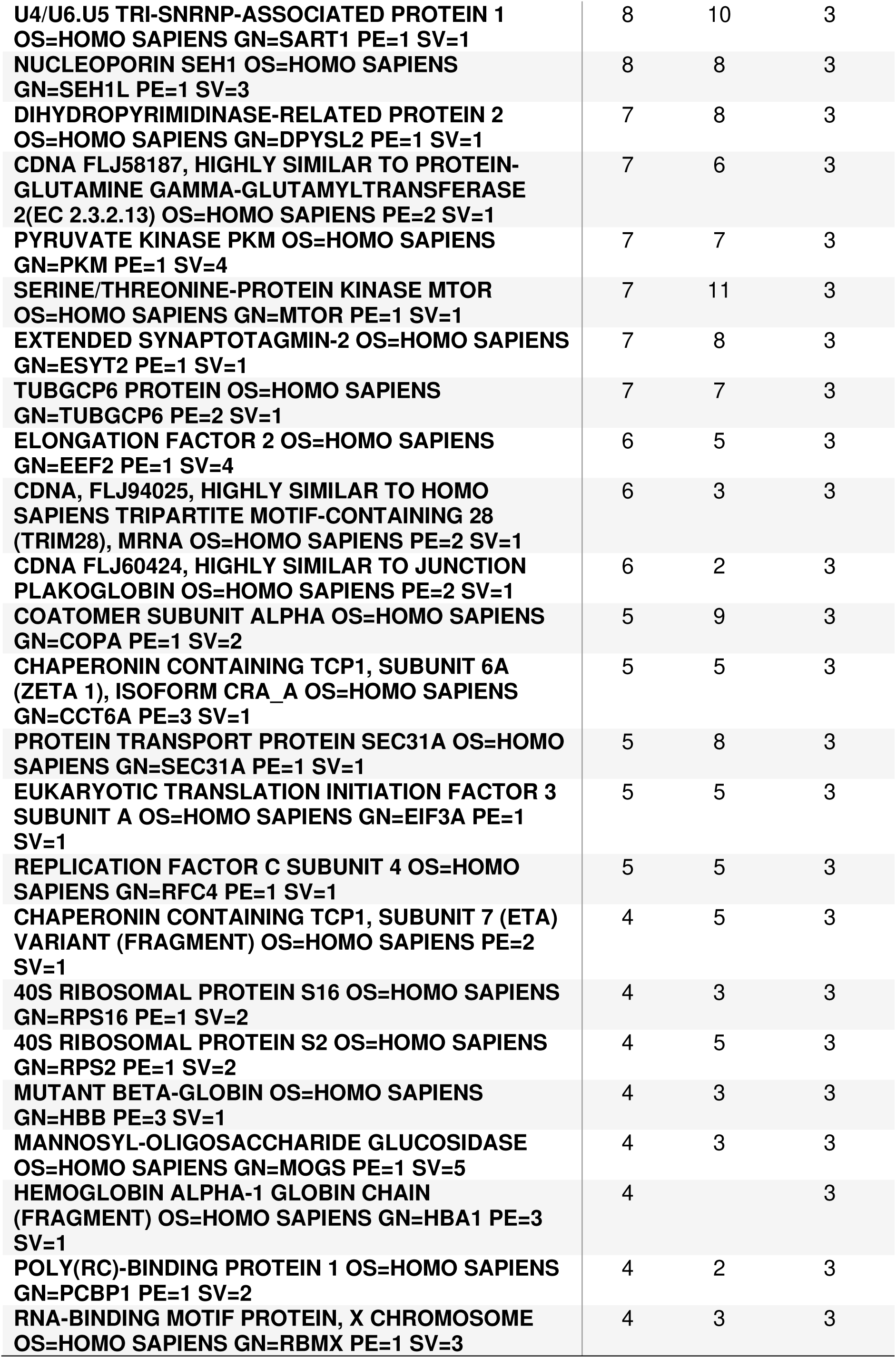
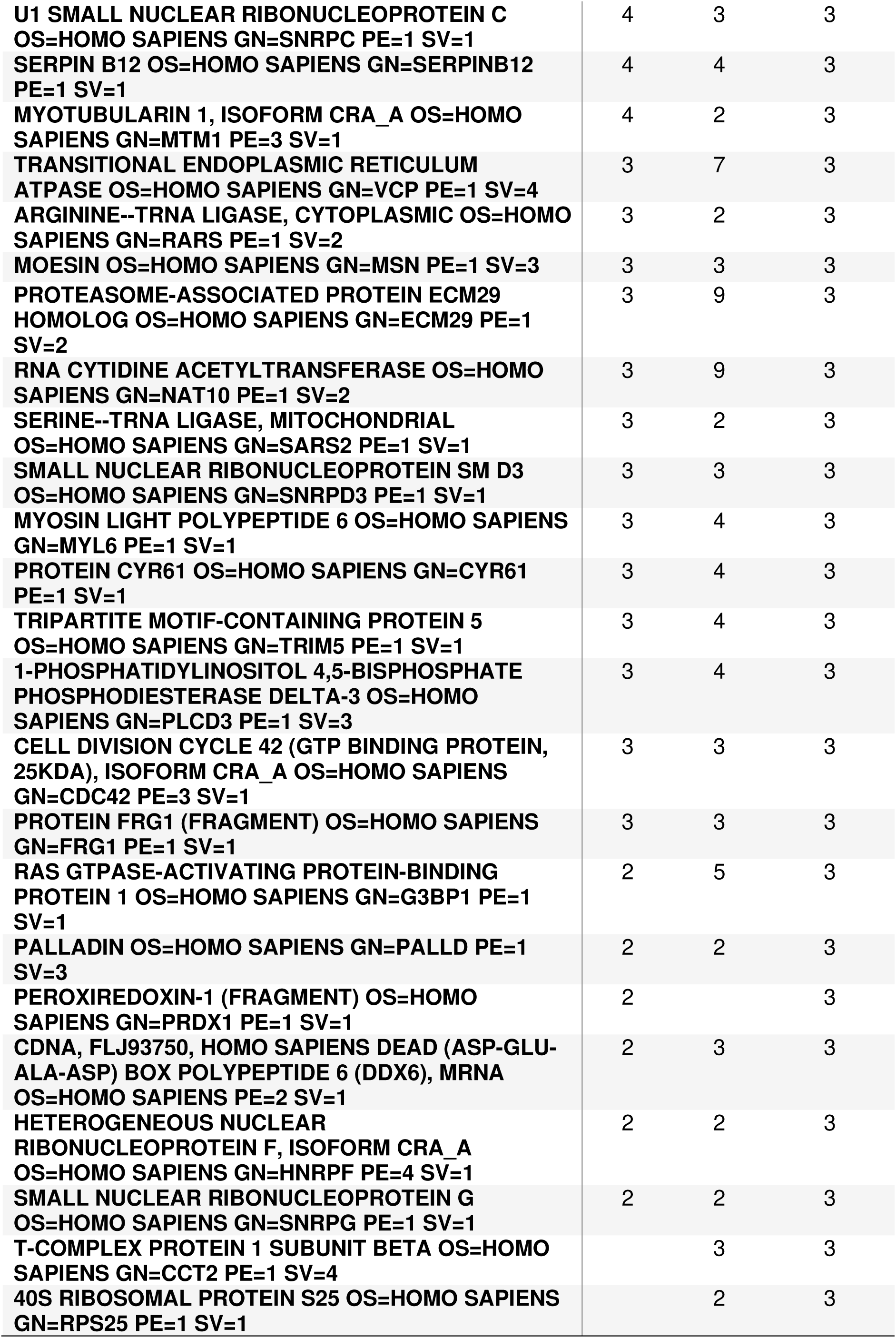
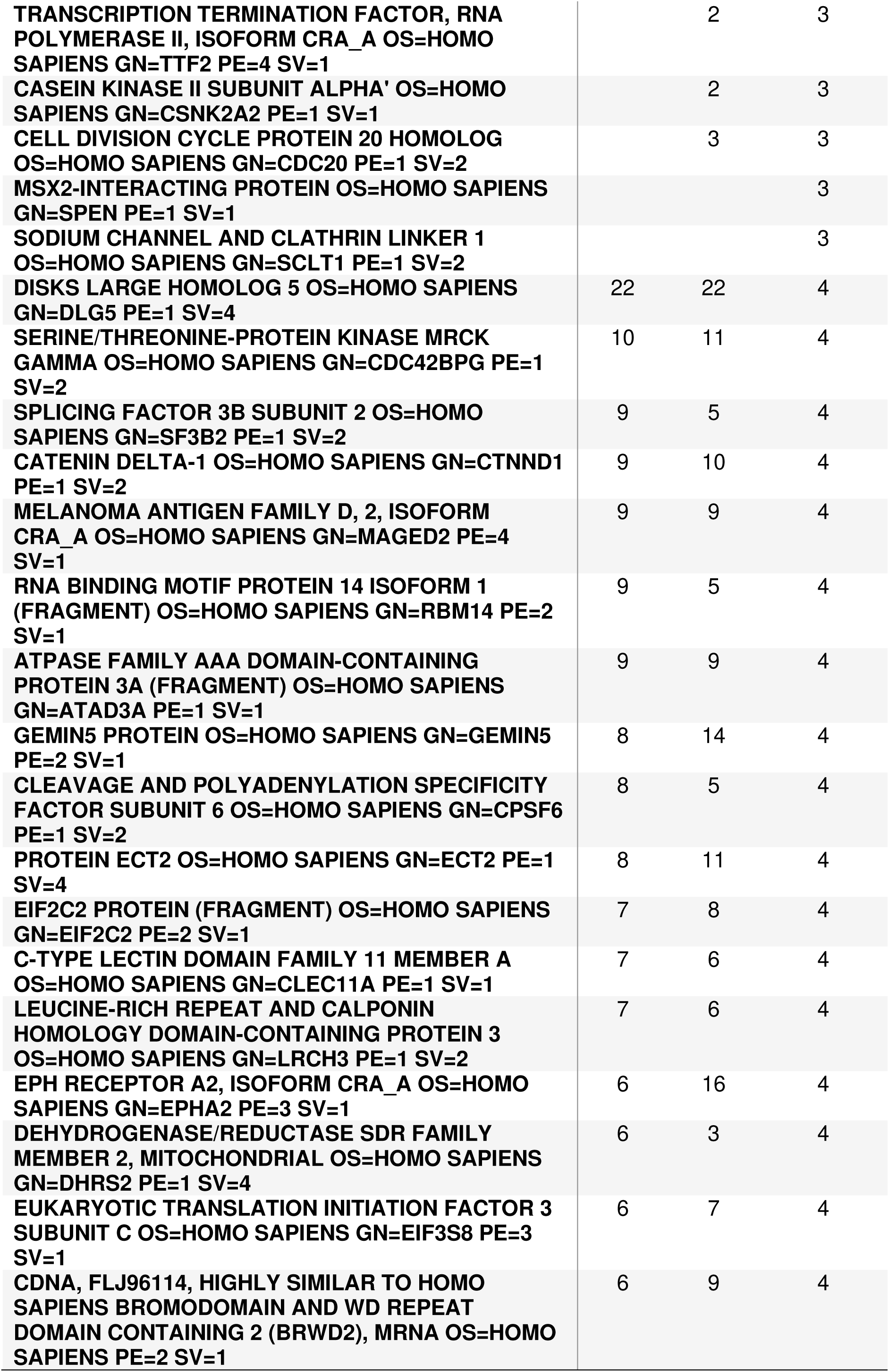
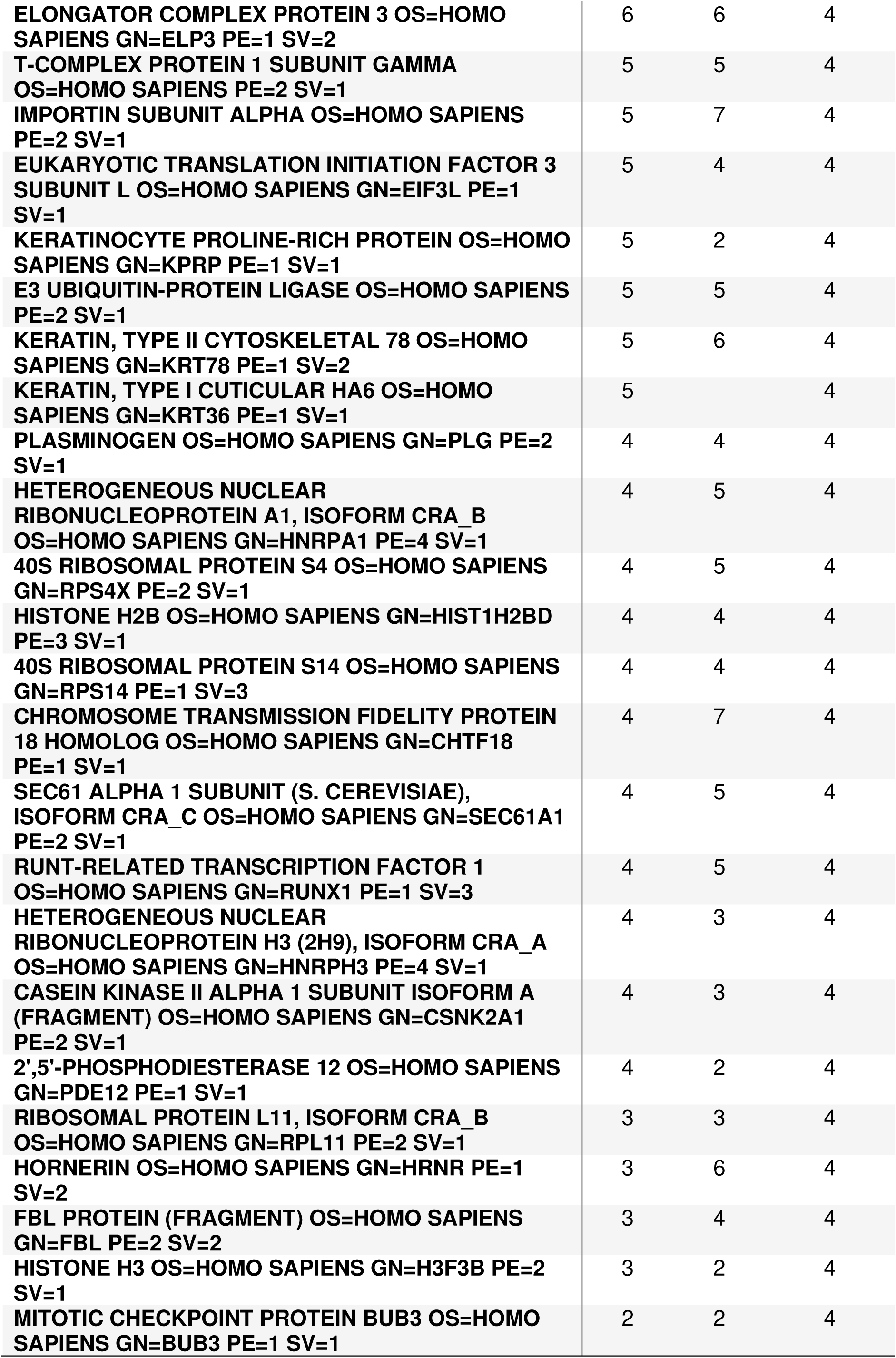
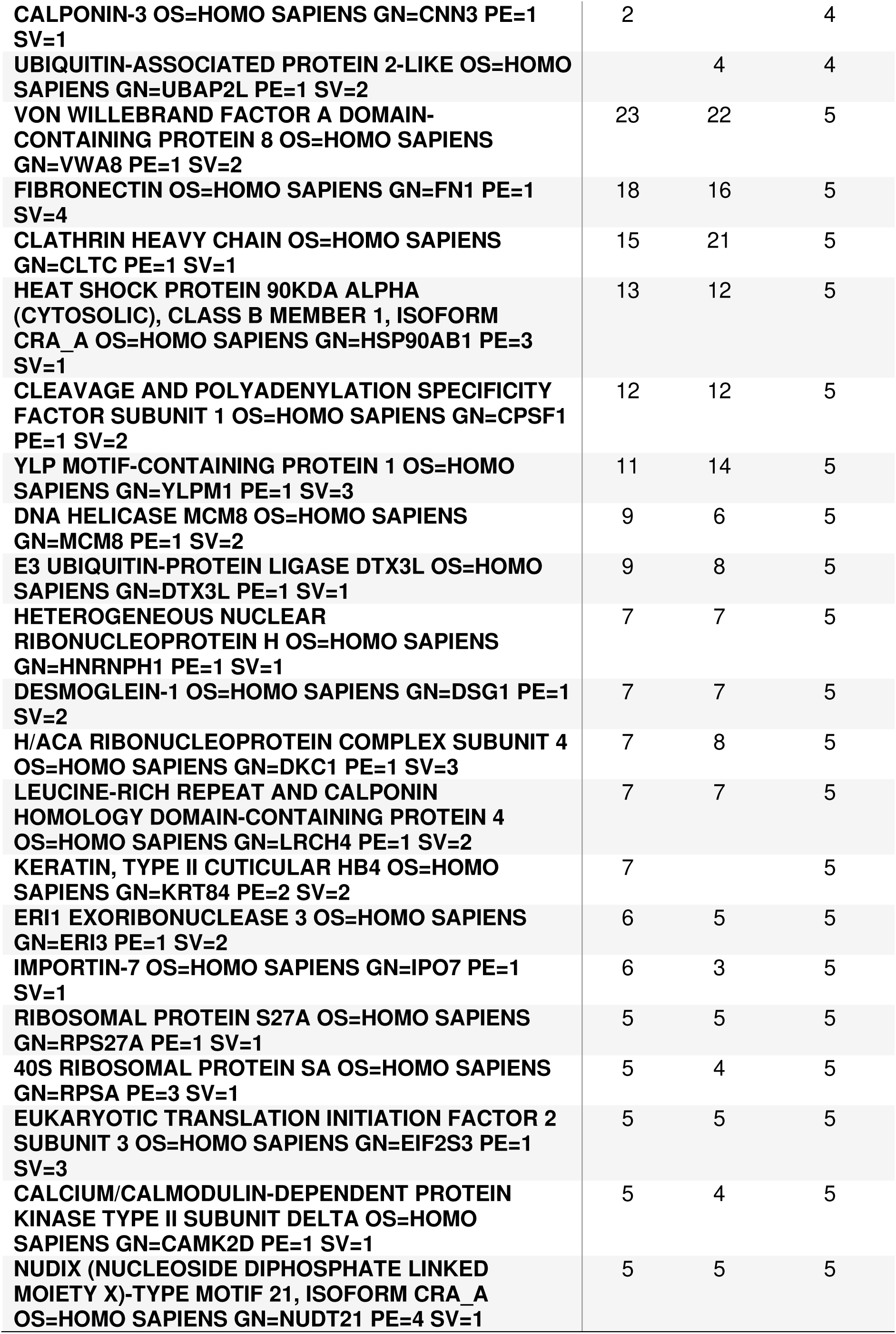
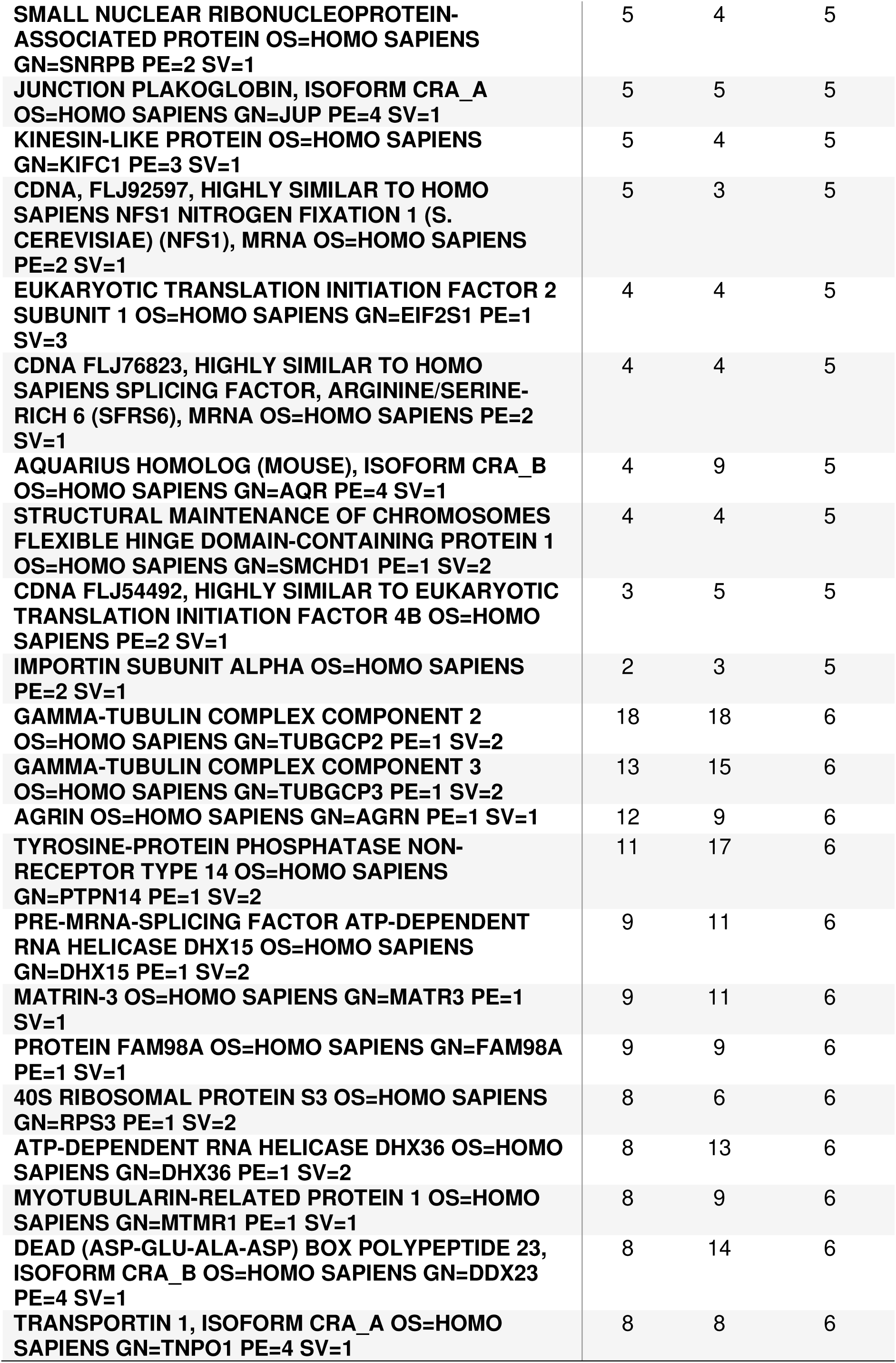
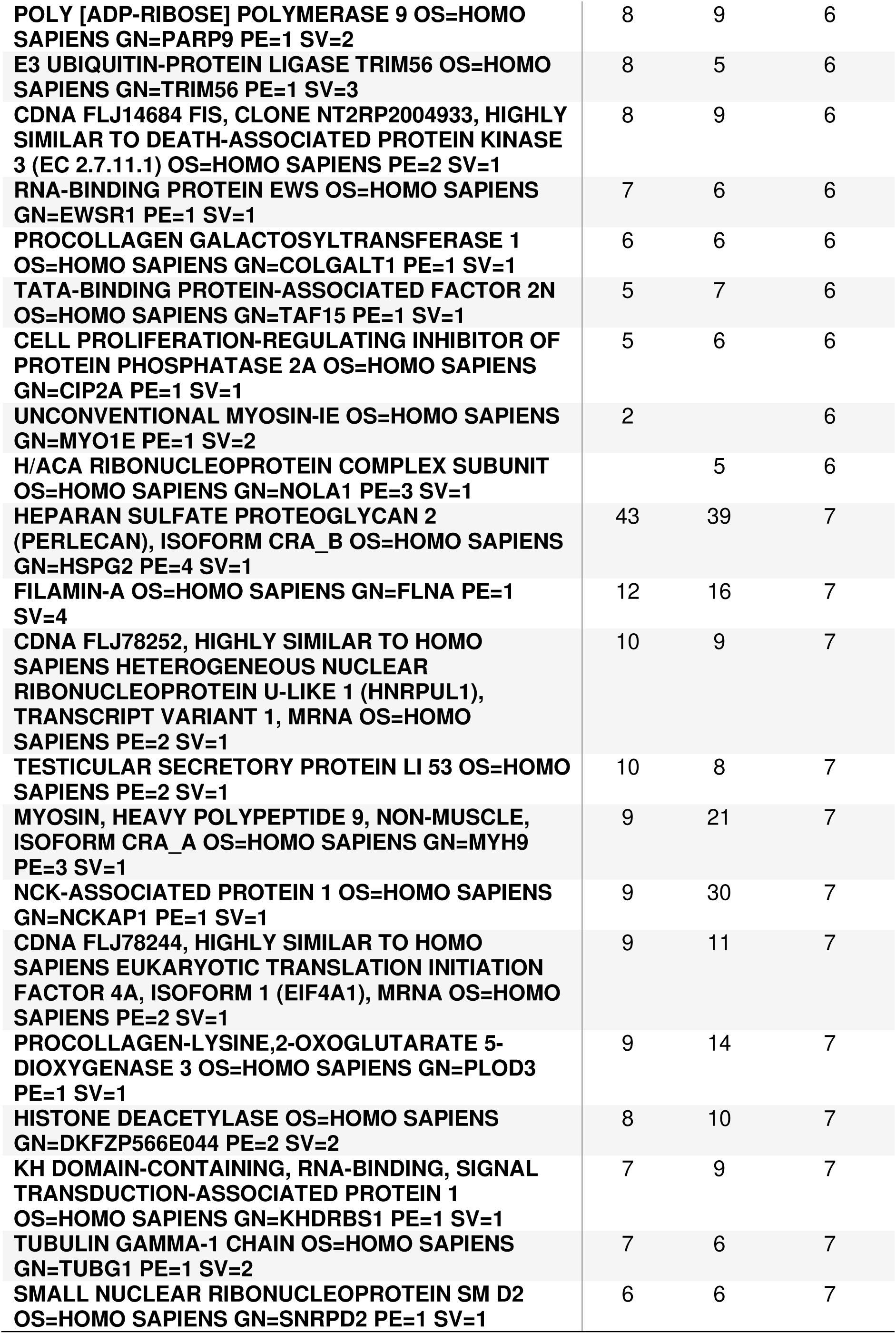
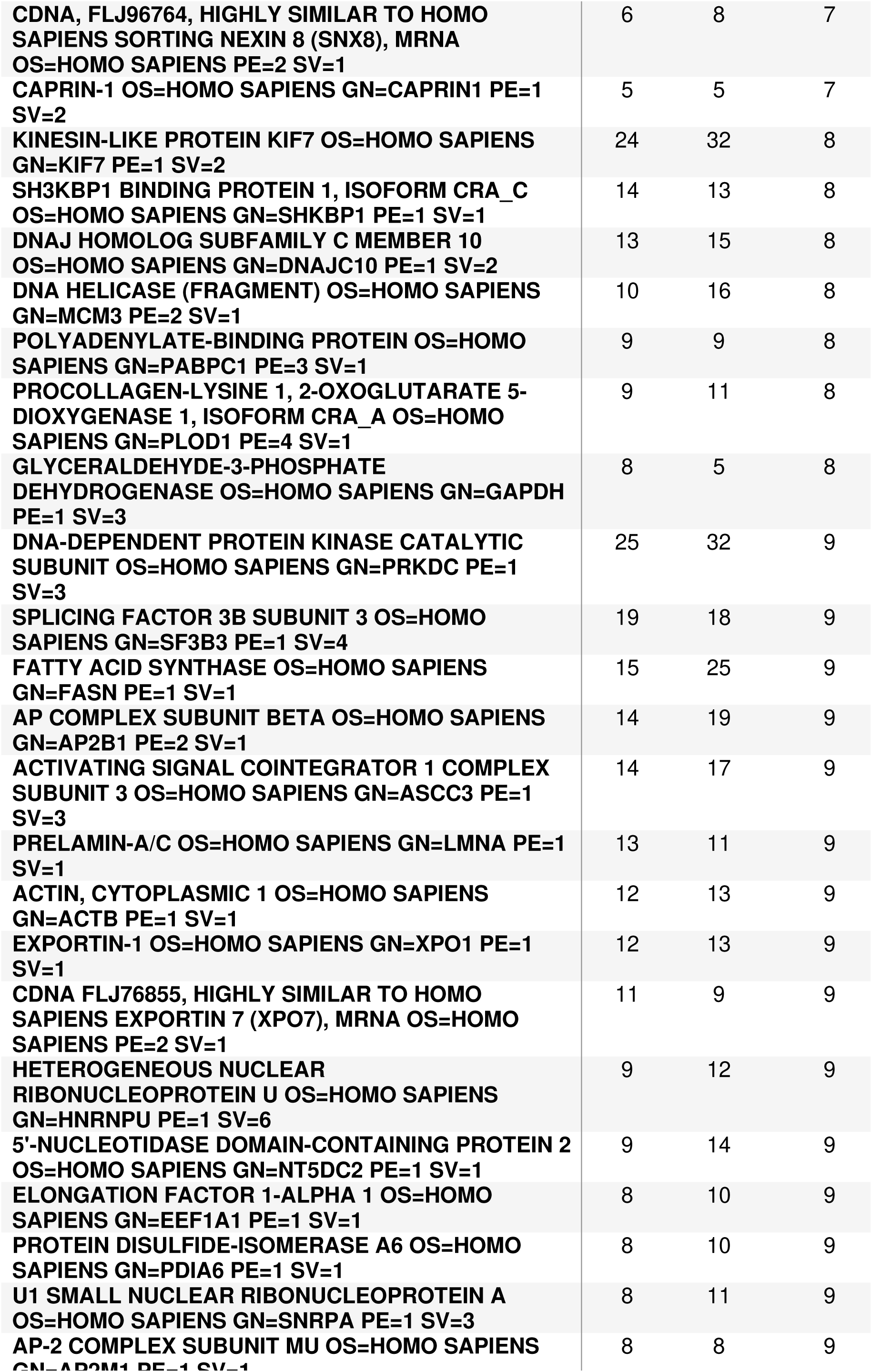
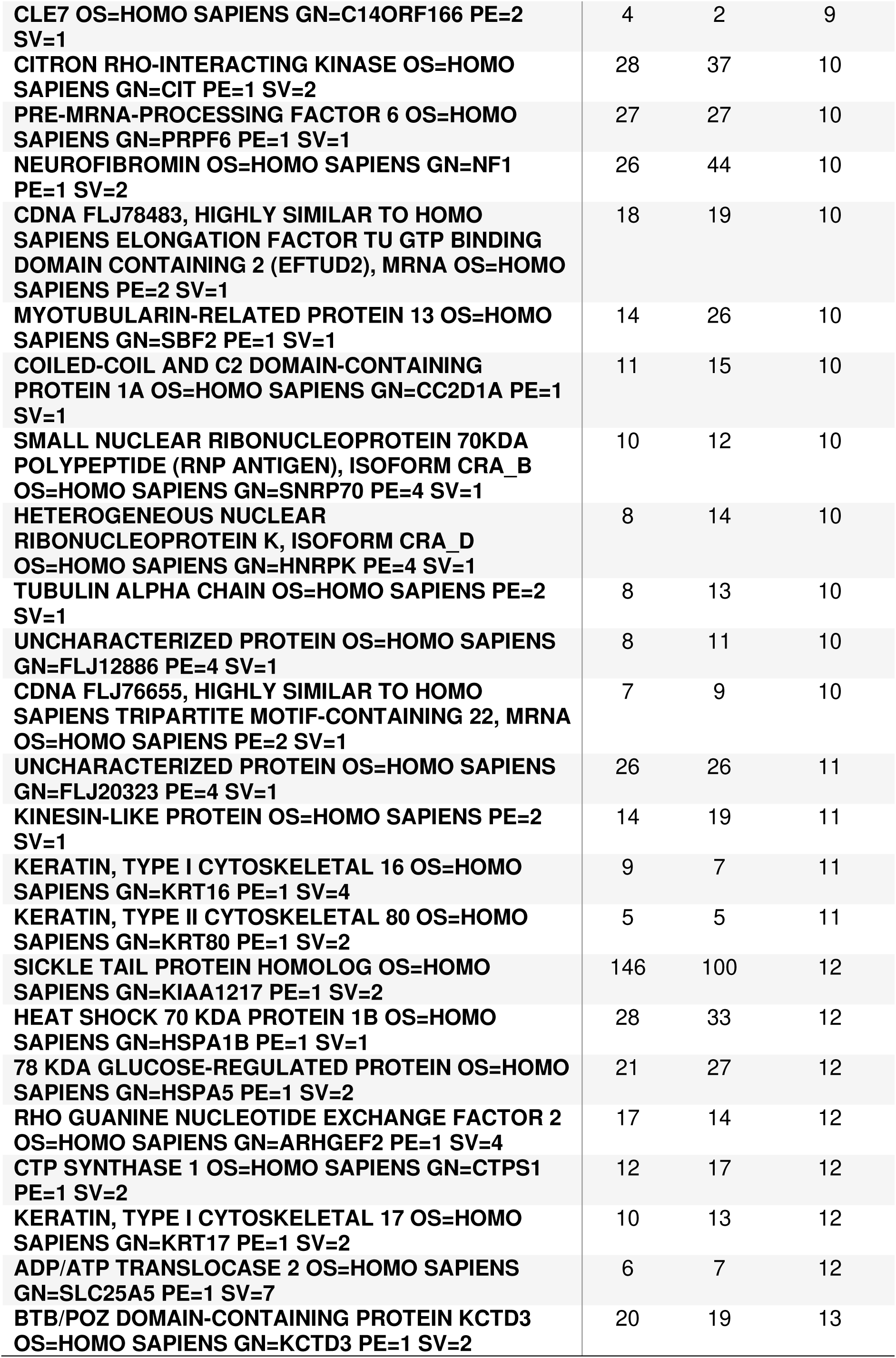
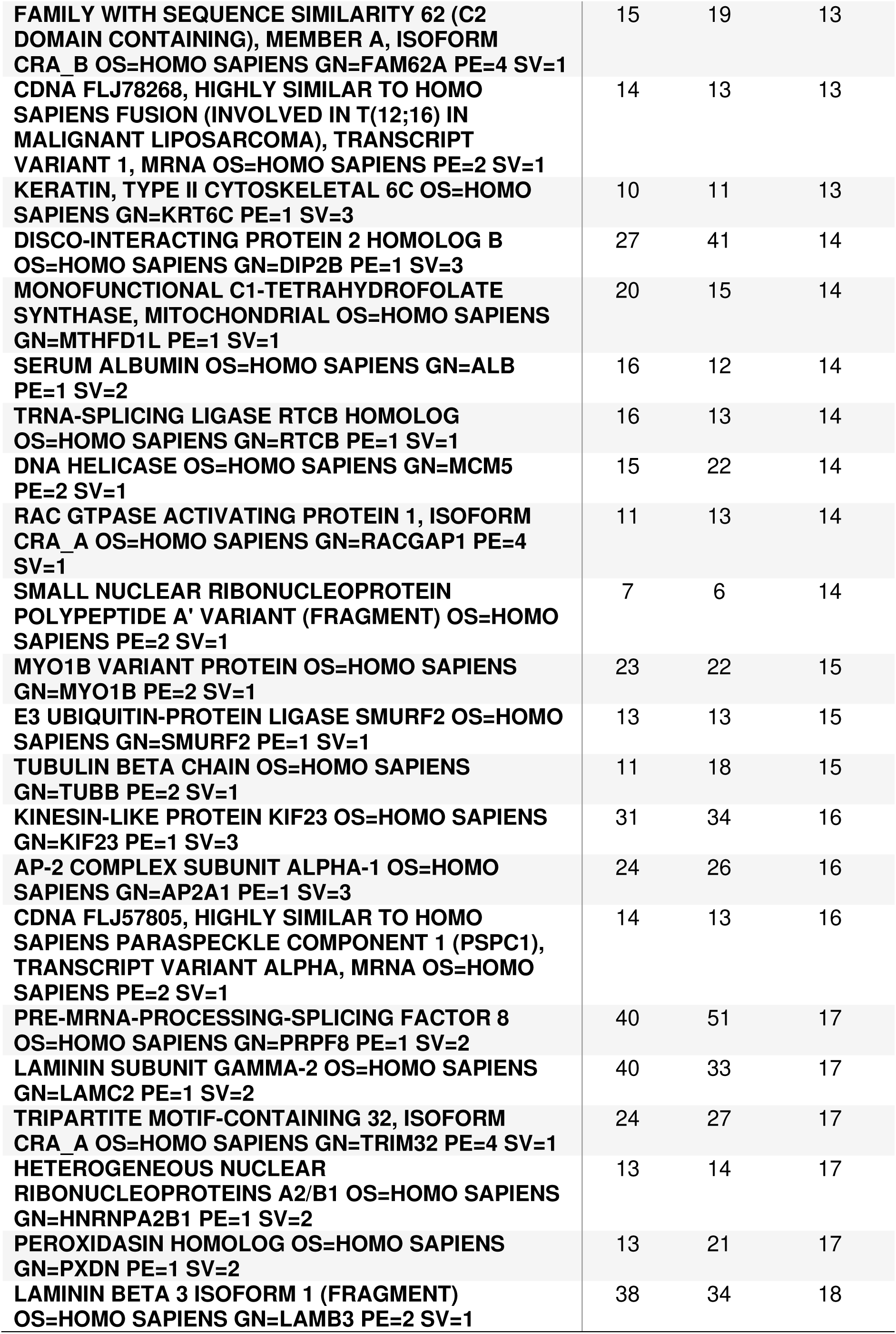
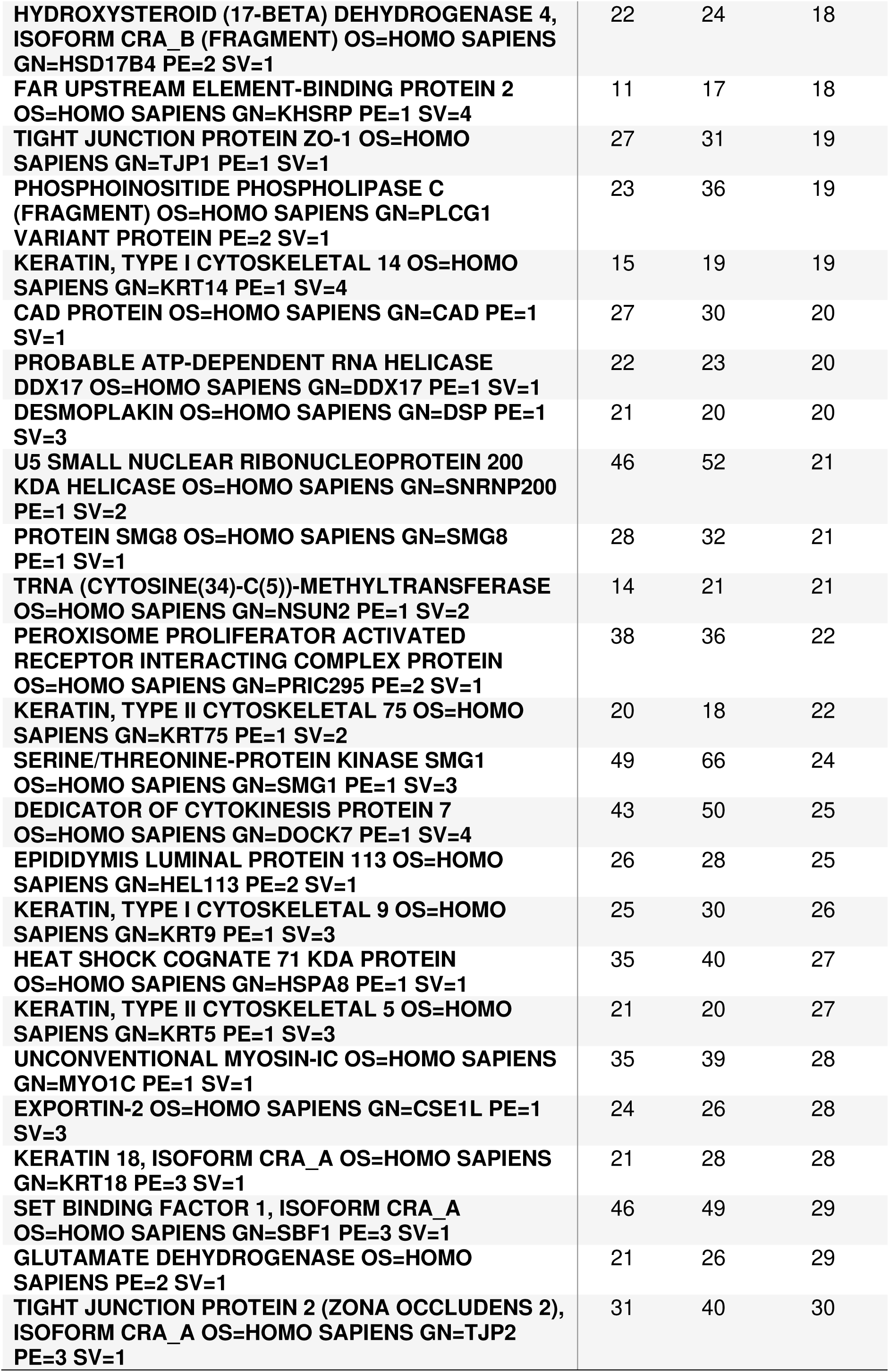
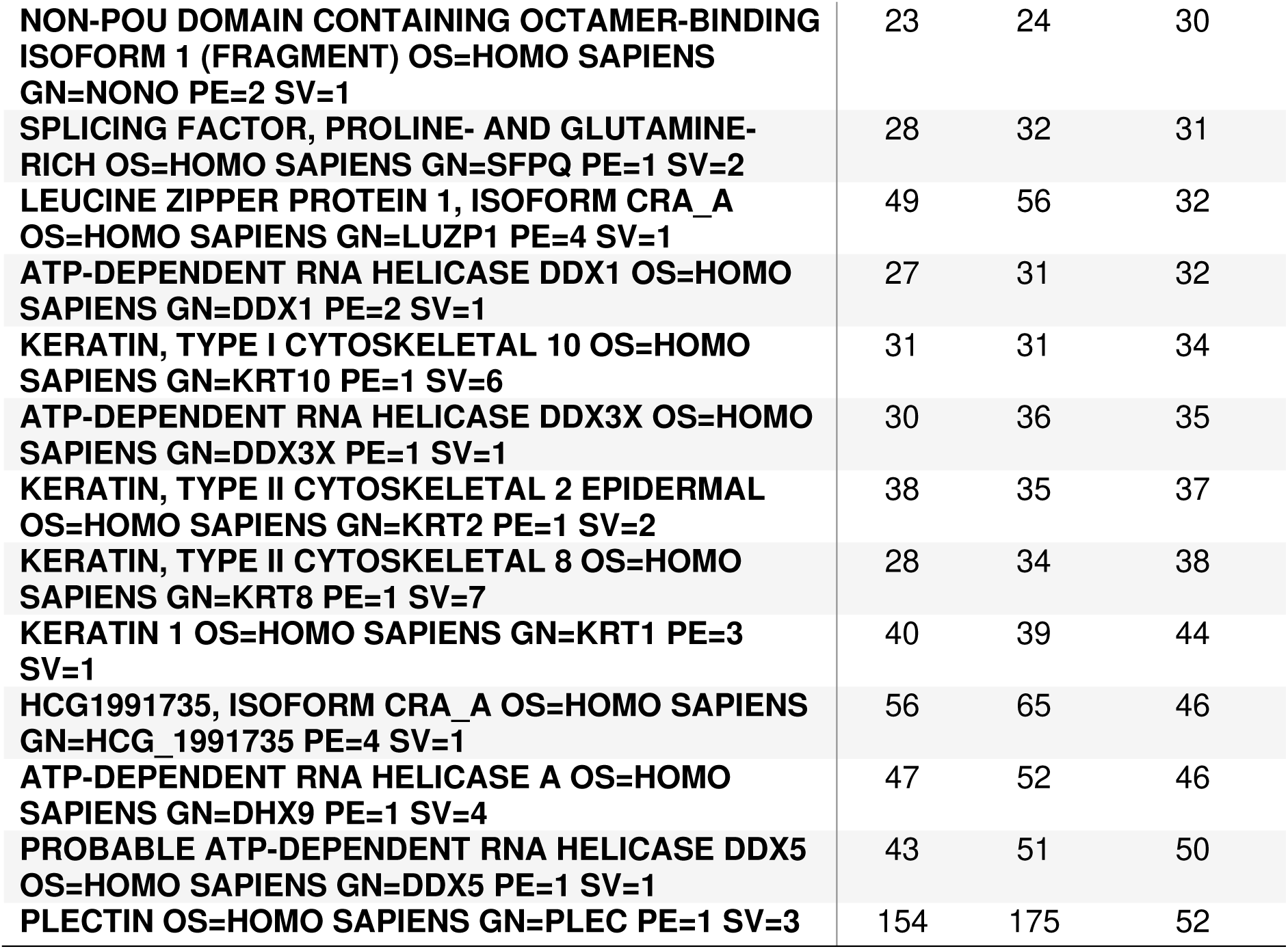
SKT interacting proteins identified with GFP-Trap immunoprecipitation and mass spectrometry analysis.

